# The Complex Simplicity of the Brittle Star Nervous System

**DOI:** 10.1101/194316

**Authors:** Olga Zueva, Maleana Khoury, Thomas Heinzeller, Daria Mashanova, Vladimir Mashanov

## Abstract

Brittle stars (Ophiuroidea, Echinodermata) have been increasingly used in studies of animal behavior, locomotion, regeneration, physiology, and bioluminescence. The success of these studies directly depends on good working knowledge of ophiuroid nervous system. Here, we describe the arm nervous system at different levels of organization: microanatomy of the radial nerve cord and peripheral nerves, neural ultrastructure, and localization of different cell types using specific antibody markers. We standardize the nomenclature of nerves and ganglia and provide an anatomically accurate digital 3D model of the arm nervous system as a reference for future studies. Our results helped identify several general features characteristic to the adult echinoderm nervous system, including the extensive anatomical interconnections between the ectoneural and hyponeural components and neuroepithelial organization of the central nervous system with its supporting scaffold formed by radial glial cells. In addition, we provide further support to the notion that the echinoderm radial glia is a complex and diverse cell population. We also tested the suitability of a range of specific cell-type markers for studies of the brittle star nervous system and established that the radial glial cells are reliably labeled by the ERG1 antibodies, whereas the best neuronal markers are acetylated tubulin, ELAV and synaptotagmin B. The transcription factor Brn1/2/4, a marker of neuronal progenitors, is expressed not only in neurons, but also in a subpopulation of radial glia. For the first time, we describe putative ophiuroid proprioceptors associated with the hyponeural part of the central nervous system.

## 1 Introduction

Brittle stars are emerging model organisms in modern biology. They have been increasingly used to address a wide range of fundamental questions, including post-traumatic regeneration of lost body appendages (Bannister et al., 2007; Purushothaman et al., 2015; Czarkwiani et al., 2016), organization and physiology of the mutable connective tissue (Wilkie, 2016), and bioluminescence (Delroisse et al., 2017). Brittle stars are also among the fastest-moving echinoderms capable of coordinated complex locomotory behaviors. The neurobiology of brittle star locomotion has been receiving attention in the contexts of body plan evolution, neurobiology, and robotics. Unlike many other members of the phylum they do not use their numerous small podia for movement. Instead, they quickly propel their body over the substratum via rapid large-scale rowing-like movements of highly motile segmented body appendages called arms. The behavior of individual five arms is centrally coordinated to produce a true bilateral movement pattern (Astley, 2012; Matsuzaka et al., 2017). To achieve this level of coordination, the nervous system must be able to coordinate muscular activity between different segments within each individual arm, as well as large-scale movements across the five arms.

All the above phenomena are controlled by or depend on the nervous system. Nevertheless, the ophiuroid nervous system has never been comprehensively studied with modern techniques. The last and only complete microanatomical description dates back to the 19th century (Hamman, 1889) and remains the best reference to date. Most modern studies report on individual aspects of echinoderm neurobiology (e.g., immunostaining with one or a few cell type markers, ultrastructure of certain regions of the nervous system). These isolated studies, although each valuable in itself, do not assemble together into general cohesive picture. The scope of this paper is, therefore, to provide a comprehensive view of the organization of the nervous system in the brittle star arm that can serve as a reference for future studies in the biology of brittle stars and echinoderms in general. We use a synthesis of different techniques to characterize various aspect of the neural architecture. These experimental approaches include three-dimensional (3D) modeling, transmission electron microscopy, and immunostaining with a series of cell-type specific glial and neuronal markers coupled with laser scanning confocal microscopy.

Here, we:

1. present an anatomically precise digital 3D model of the nervous system in an arm segment. We made an effort to trace the origin and targets of all major peripheral nerves and propose standardized terminology for them.
2. provide a detailed description of the neurohistological organization of both the radial nerve cord and peripheral nerves and ganglia and discuss both the features that ophiuroids share with other echinoderms and unique ophiuroid characteristics
3. describe the immunoreactivity of the cells of the nervous tissue with cell type-specific antibody markers.

Together, our data help establish both the general principles of neural architecture common to the phylum Echinodermata and the specific ophiuroid features. We also confirmed and expanded earlier observations (Mashanov et al., 2015a) of complex and molecularly heterogeneous organization of echinoderm glia. This study also describes for the first time putative proprioceptors embedded in the CNS.

## 2 Materials and Methods

### 2.1 Animal collection and maintenance

Adults of *Amphipholis kochii* Lütken, 1872 were collected from Vostok Bay, Sea of Japan (Russia). Adult individuals of *Ophioderma brevispinum* Say, 1825 were purchased from Gulf Specimen Marine Laboratories, Inc. (Panacea, FL). The animals were kept in glass aquaria with aerated sea water.

### 2.2 Electron microscopy

For transmission electron microscopy (TEM), arms of *A. kochii* were fixed in 2.5% glutaraldehyde dissolved in 0.05 M cacodylate buffer (pH 7.6) for 24 h at 4°C. After fixation, the specimens were rinsed in the same buffer and postfixed in 1% OsO_4_ in cacodylate buffer for 1 h. The tissue samples were then decalcified in several changes of a solution containing 1% ascorbic acid and 0.15 M NaCl, dehydrated in a graded series of ethanol and acetone and embedded in the Araldite epoxy resin. Sections were cut with glass knives on Ultracut E (Reichert, Vienna, Austria) and UC6 (Leica) ultratomes. Ultrathin (50 – 70 nm) sections were stained with aqueous uranyl acetate and lead citrate and then examined and photographed with a Zeiss EM 10 transmission electron microscope.

For scanning electron microscopy (SEM), specimens were fixed in 2.5% glutaraldehyde in 0.05 M cacodylate buffer at pH 7.6, dehydrated in ethanol followed by an acetone series, critical point dried, and then sputter coated with carbon and gold. Specimens were examined with a Jeol JSM-IC848 scanning electron microscope (JEOL Ltd., Tokyo, Japan).

### 2.3 3D surface reconstruction

Complete series of transverse semi-thin (0.8 *μ*m) sections were cut from one arm segment. For this purpose, we used one of the Araldite-embedded tissue samples that were prepared for TEM (see above). The sections were collected on gelatin-coated slides, stained with 1% toluidine blue in 1% aqueous sodium borate and mounted in DPX (Fluka). Every sixth section in the series was photographed with a Jenamed 2 (Carl Zeiss Jena) light microscope equipped with a Leica DC 150 digital camera. Preliminary image processing (brightness and contrast adjustments, etc.) were performed using Adobe Photoshop CS2 software. The stack of digitized micrographs was then imported into the Amira 3.1.1 volume modeling and visualization software (Mercury Computer Systems, Inc., Chelmsford, MA, USA), which was used for stack alignment, segmentation, and generation of the initial 3D model. Final editing of the model was performed in Blender, an open-source 3D editor (https://www.blender.org). Rendered images were generated using Blender’s *Cycles* engine with 500 samples.

The original stack of images, saved as a video file, is available in Supplementary File 1. The final 3D model in various interactive and non-interactive (video) formats can be accessed in Supplementary Files 2 thru 11.

### 2.4 Immunohistochemistry

Fluorescent immunohistochemistry on frozen sections was performed as described elsewhere (Mashanov et al., 2013). Briefly, animals were anesthetized in 0.2% chlorobutanol (Sigma) and then portions of the arm approximately 5 segments long were fixed in 4% paraformaldehyde prepared in 0.01 M PBS (pH 7.4) overnight at 4°C. For immunostaining with anti-synaptotagmin antibodies, the samples were fixed in ice-cold methanol for 30 min. The samples were then washed in PBS, decalcified in 10% EDTA, cryoprotected in sucrose and embedded in the Tissue-Tek O.C.T. Compound (Sakura). Cryosections (10 *μ*m thick) were cut using Leica CM1860 cryostat, collected on gelatin-covered slides and incubated at 42°C overnight. The slides were then washed in PBS incubated in 0.1 M glycine for 1 h to quench autofluorescence. After another wash in PBS (3×10 min), the sections were blocked using 2% goat serum for 1 h. The first antibodies (see Table 1) were applied at 4°C overnight. After extensive washing in PBS (4×10 min), the sections were incubated in the secondary antibodies (Table 1) for 1 h at room temperature. Unbound antibodies were removed in four changes of PBS (10 min each) and nuclei were stained with 5 *μ*M DRAQ5 (Thermo Scientific) for 30 min. After the final round of washes (3×5 min), the slides were coverslipped in an anti-fading medium containing 2.5% DABCO and 10% Mowiol 4-88 in 25% buffered glycerol (0.2M Tris-HCL, pH 8.5).

**Table 1:** Antibodies used in this study

| Antibodies used | Source | Host species | Dilution |
| --- | --- | --- | --- |
| Acetylated tubulin | Sigma (T6793) | Mouse | 1:1,000 |
| Brn1/2/4 | Garner et al. (2016) | Rat | 1:1,000 |
| ELAV | Garner et al. (2016) | Rabbit | 1:3,000 |
| ERG1 | Mashanov et al. (2010) | Mouse | 1:1 |
| GFSKLYFamide | Díaz-Miranda et al. (1995) | Rabbit | 1:1,000-1:5,000 |
| Goat anti-mouse Cy3 | Jackson ImmunoResearch (115-165-146) | Goat | 1:2,000 |
| Goat anti-rabbit Cy3 | Jackson ImmunoResearch (115-165-144) | Goat | 1:1,000 |
| Goat anti-rabbit FITC | ThermoFisher Scientific (65-6111) | Goat | 1:200 |
| Goat anti-rat FITC | GenWay (GWB-7B4D70) | Goat | 1:100 |
| Synaptotagmin B | Nakajima et al. (2004) | Rat | 1:5,000 |

Whole-mount staining was performed in a similar way with the following modifications. After the fixation and decalcification steps, the tissues were bleached in increasing concentrations of hydrogen peroxide (0.3%, 1%, and 3%) and then permeabilized by Proteinase K digestion (2.5 *μ*g/ml, 15 min at room temperature). All wash buffers contained 0.5% Triton X-100. The incubation steps in the first and second antibodies were increased to 2 days each and performed at 4°C.

Stacks of optical sections were taken using the Olympus confocal laser scanning microscope FV1000. Maximum intensity Z-projections were generated in the Fiji image processing software (Schindelin et al., 2012).

Unless indicated otherwise, images of whole-mount specimens and micrographs of longitudinal sections are oriented with the distal side to the right.

## 3 Results

### 3.1 Anatomical organization of the nervous system in the arm

#### Nerve Ring and Radial Nerve Cord

The brittle star body is composed of a flattened disk and five long unbranched segmented arms (Fig 1A, B). Each segment, contains a set of skeletal elements, including the central vertebral calcareous ossicle surrounded by four peripheral arm shields or plates: a dorsal, a ventral, and two lateral ones (Fig 1D, D’). Each lateral arm shield bears a vertical row of arm spines (Fig 1B). On the ventral surfaces of the arm segment, a pair of podia emerges through pores adjacent to bases of the ventral arm spines. Each podium is protected by two tentacle scales (Fig 1B). The vertebrae of adjacent segments are joined together by paired aboral and oral intervertebral muscles and the intervertebral ligament that has a complex geometry (Fig 1C–D’; 3C, E).

The main components of the central nervous system in brittle stars are organized into a pentaradial pattern and thus correspond to the overall layout of the general body plan. In each arm, the radial nerve cord (RNC) lies beneath the water-vascular canal (Fig 1C–D’; 3A, E) beneath the oral plate and ligament. At the attachment of the arm to the disk, each of the the RNC approaches the centrally located esophagus, ascends aborally and bifurcates (Fig 2). The side branches of adjacent RNCs fuse together to form a continuous nerve ring Fig 2B). Both the RNCs and the nerve ring are composed of two layers of nervous tissue, a thicker orally located ectoneural part and a much thinner hyponeural tissue that covers the aboral surface of the ectoneural cords (Fig 3, 4A).

**Figure 1:**
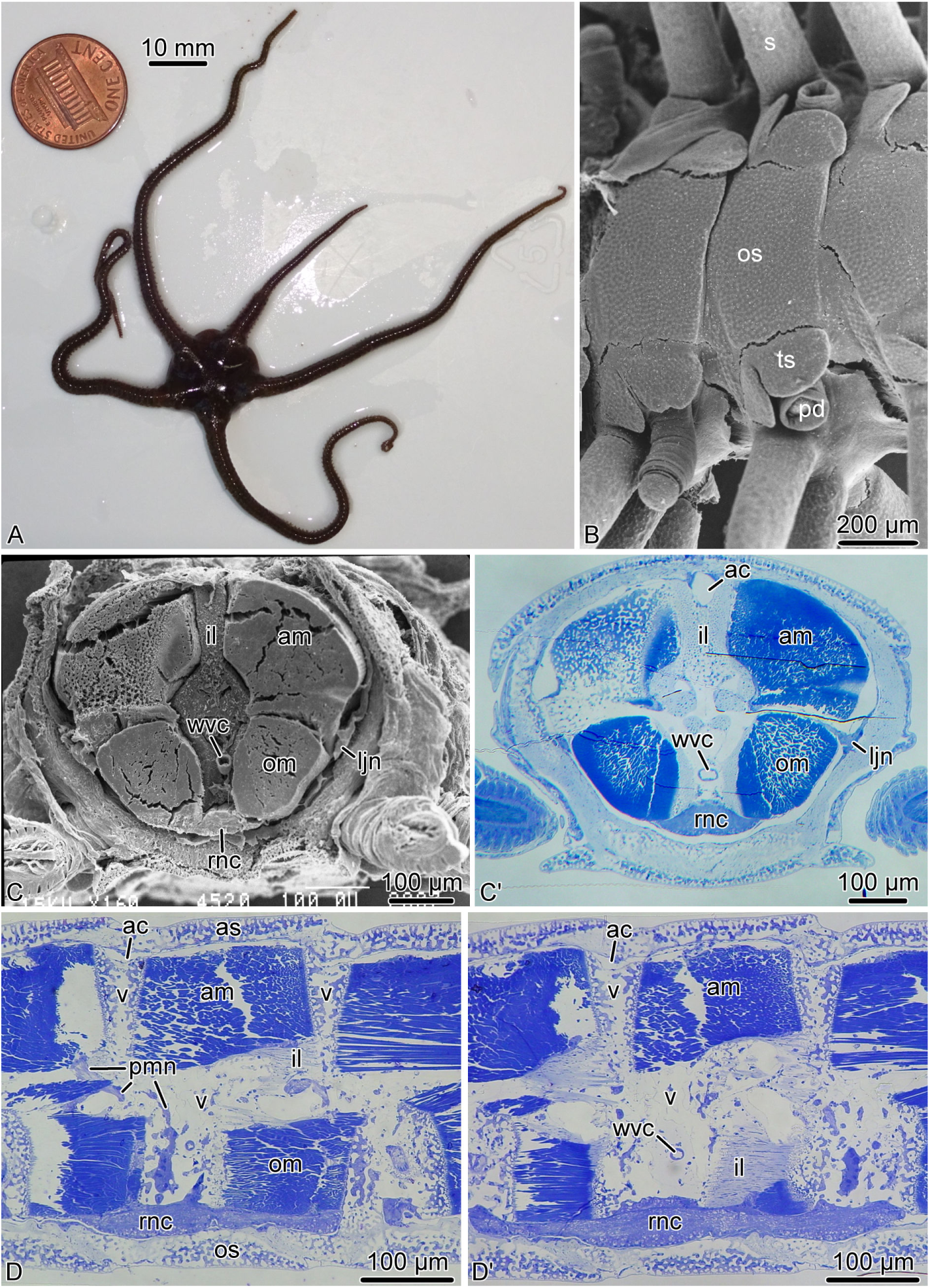
Brittle star arm anatomy. **A.** An individual of *O. brevispinum*. **B.** Scanning electron micrograph of the oral arm surface of *A. kochii*. C and C’. Cross section of the arm of *A. kochii*. **C.** The arm was embedded into epoxy resin and sectioned transversely. The resin was then removed, and the cut surface was imaged with a scanning electron microscope. C’. The corresponding plastic section stained with methylene blue. D and D’. Longitudinal paragsagittal sections through the arm in *A. kochii* stained with methylene blue. The skeletal elements are not visible in any of the micrographs, because they were removed during tissue processing. Abbreviations: *ac* – arm coelom; *am* – aboral intervertebral muscle; *as* – aboral shield; *il* – intervertebral ligament; *ljn* – lateral juxtaligamental node; *om* – oral intervertebral muscle; *os* – oral shield; *pd* – podium; *pmn* – hyponeural proximal muscle nerve; *rnc* – radial nerve cord; *s* – spine; *ts* – tentacle scale; *wvc* – water-vascular canal; *v* – vertebral ossicle.

**Figure 2:**
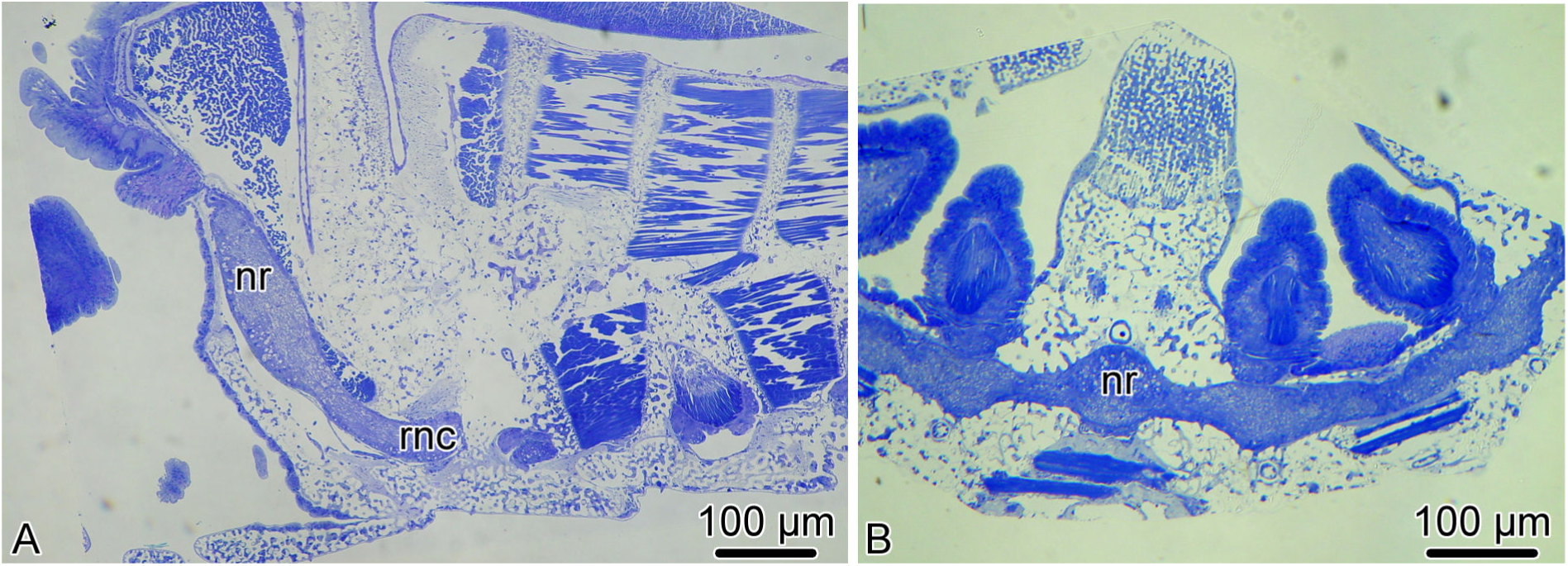
Transition between the radial nerve cord and the circumoral nerve ring in *A. kochii*. Semi-thin sections stained with methylene blue. **A.** Radial section (parallel to the long axis of the arm). **B.** Horizontal section (i.e., orthogonal with respect to the oral-aboral axis) through the nerve ring. Abbreviations: *nr* – nerve ring; *rnc* – radial nerve cord.

**Figure 3:**
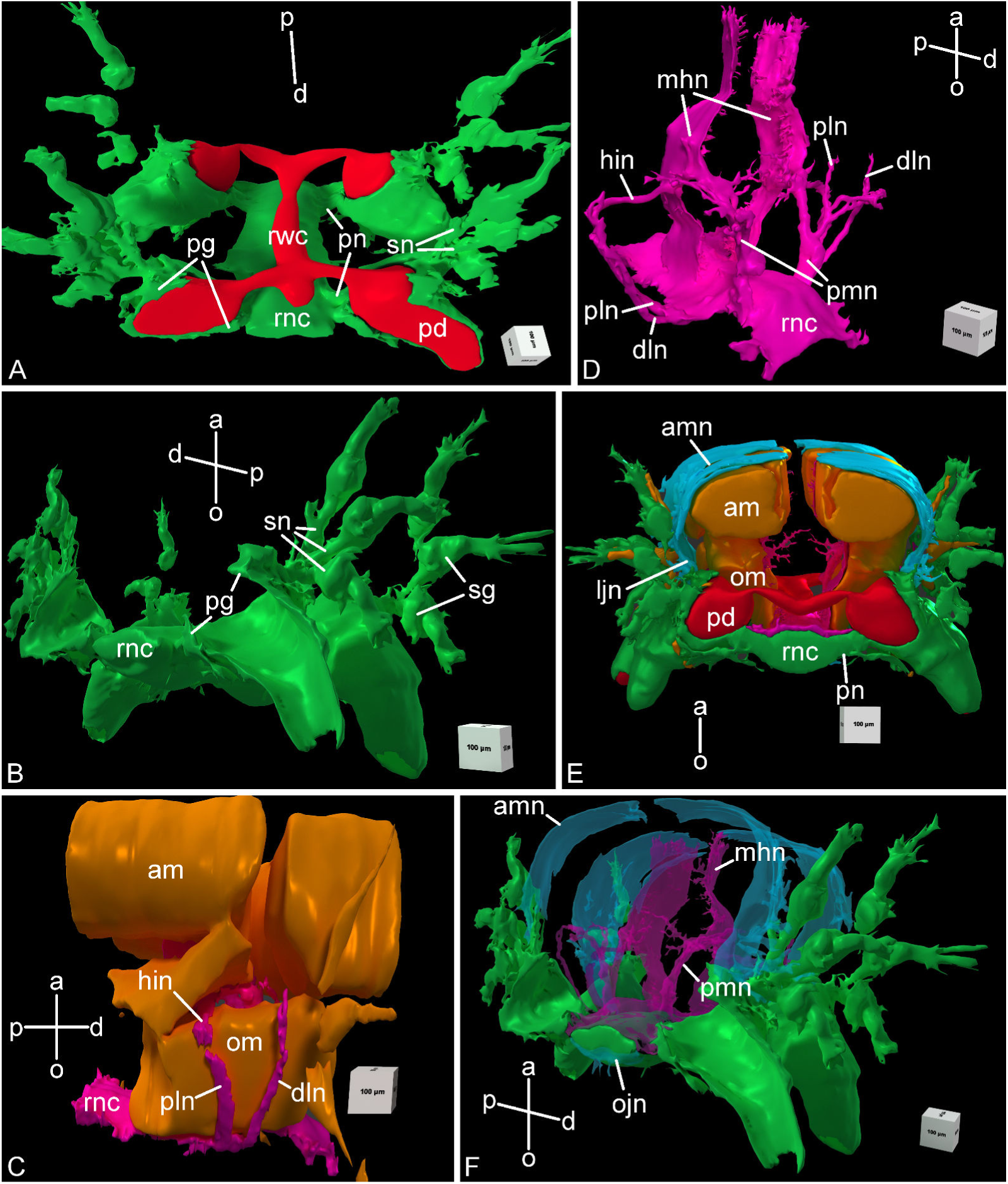
Three-dimensional reconstruction of the radial nerve cord and peripheral nerves in the arm segment in *A. kochii*. Other anatomical structures are provided for reference, when indicated. **A.** Aboral view of ectoneural system *(green)*. Components of the water-vascular system are shown in *red*. **B.** Distal view of the ectoneural system. **C.** Side view of muscles *(brown)* and the hyponeural system *(magenta)*. **D.** Oblique side the view of hyponeural system *(magenta)*. **E.** Proximal view of the complete nervous system. The muscles *(brown)* and the water-vascular system *(red)* are also shown. F. Distal view of the complete nervous system. The orientation of the projections is indicated by axes on each image: *a* – aboral; *d* – distal; *o* – oral; *p* – proximal. Abbreviations: *am* – aboral intervertebral muscle; *amn*– aboral mixed nerve; *dln* – distal lateral hyponeural nerve; *hin* – horizontal intermuscular hyponeural nerve; *ljn* – lateral juxtaligametal node; *mhn* – median hyponeural nerve; *ojn*– oral juxtaligamental node; *om* – oral intervertebral muscle; *pd* – podium; *pg* – podial ganglion; *pln* – proximal lateral hyponeural nerve; *pmn* – proximal muscle nerve; *pn* – podial nerve; *rnc* – radial nerve cord; *rwc* – radial water-vascular canal; *sg* – spine ganglion; *sn* – spine nerve.

**Figure 4:**
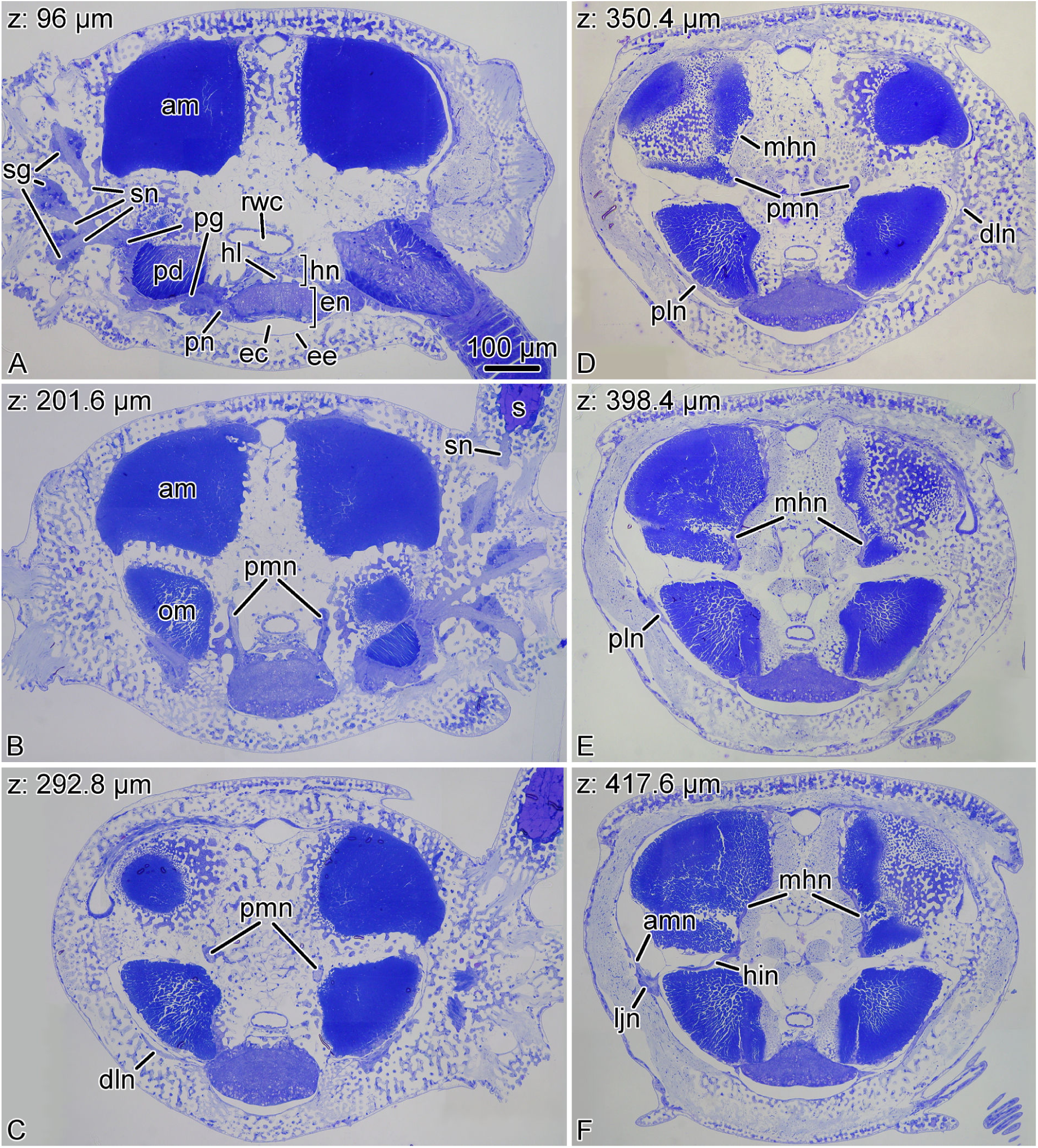
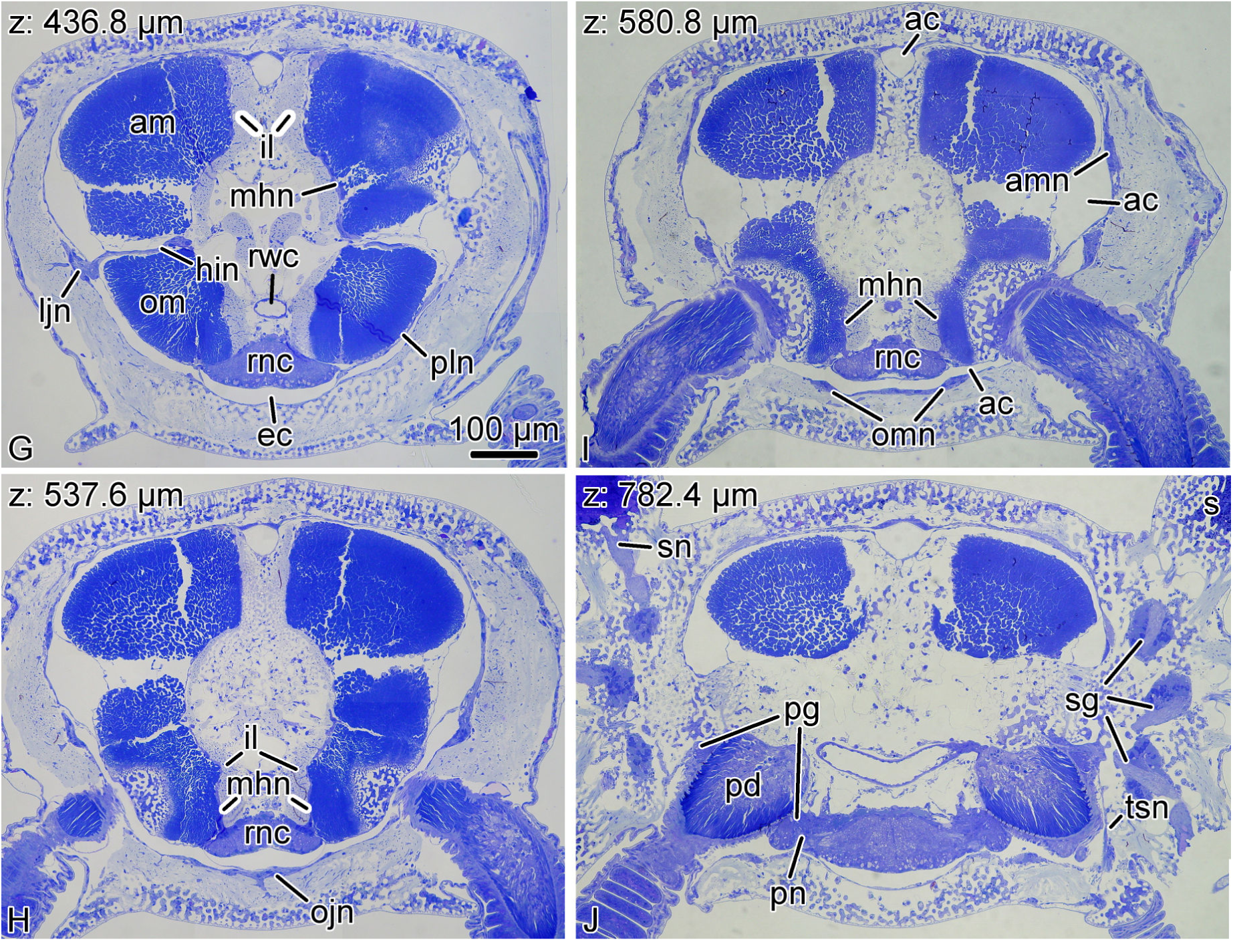
Representative semithin sections from the series that was used to generate the 3D models shown in Fig. 3. The series was cut in the proximal-to-distal direction. The Z-coordinate (the position of the section in the series) is indicated in the top left corner. Abbreviations: *ac* – arm coelom; *am* – aboral intervertebral muscle; *amn* – aboral mixed nerve; *dln* – distal lateral hyponeural nerve; *ec* – epineural canal; *ee* – epineural epithelium; *en* – ectoneural part of the radial nerve cord; *hin* – horizontal intermuscular hyponeural nerve; *hl* – hemal lacuna; *hn* – hyponeural part of the radial nerve cord; *il* – intervertebral ligament; *ljn* – lateral juxtaligametal node; *mhn* – median hyponeural nerve; *ojn* – oral juxtaligamental node; *om* – oral intervertebral muscle; *omn* – oral mixed nerve; *pd* – podium; *pg* – podial ganglion; *pln* – proximal lateral hyponeural nerve; *pmn* – proximal muscle nerve; *pn* – podial nerve; *rnc* – radial nerve cord; *rwc* – radial water-vascular canal; *s* – spine; *sg* – spine ganglion; *sn* – spine nerves; *tsn* – tentacle scale nerve.

The radial nerve cord in brittle stars has clear metameric organization and corresponds to segmented organization of other components of the arm, including the skeleton, ligaments, muscles, and water-vascular system. At the level of each vertebral ossicle, both the ectoneural and hyponeural layers of the RNC are thickened to form ganglionic swellings (Fig. 1D, D’). In each segment, the radial nerve cords gives off a complex system of peripheral nerves that innervate different metameric anatomical structures of the arm (Fig. 3). This peripheral nervous system is described below.

As in the previously studied CNS of sea cucumbers (Mashanov et al., 2006, 2016), the ectoneural and hyponeural components of the RNC have organization of tubular cords. The aboral wall of the ectoneural cord is very thick and is formed by a tall ectoneural epithelium (Fig. 4A; 5A). The opposite oral wall is composed of a very thin epineural epithelium. The cavity that separates the ectoneural neuroepithelium and the epineural epithelium constitutes the epineural canal (Fig. 5A; 6A, B).

**Figure 5:**
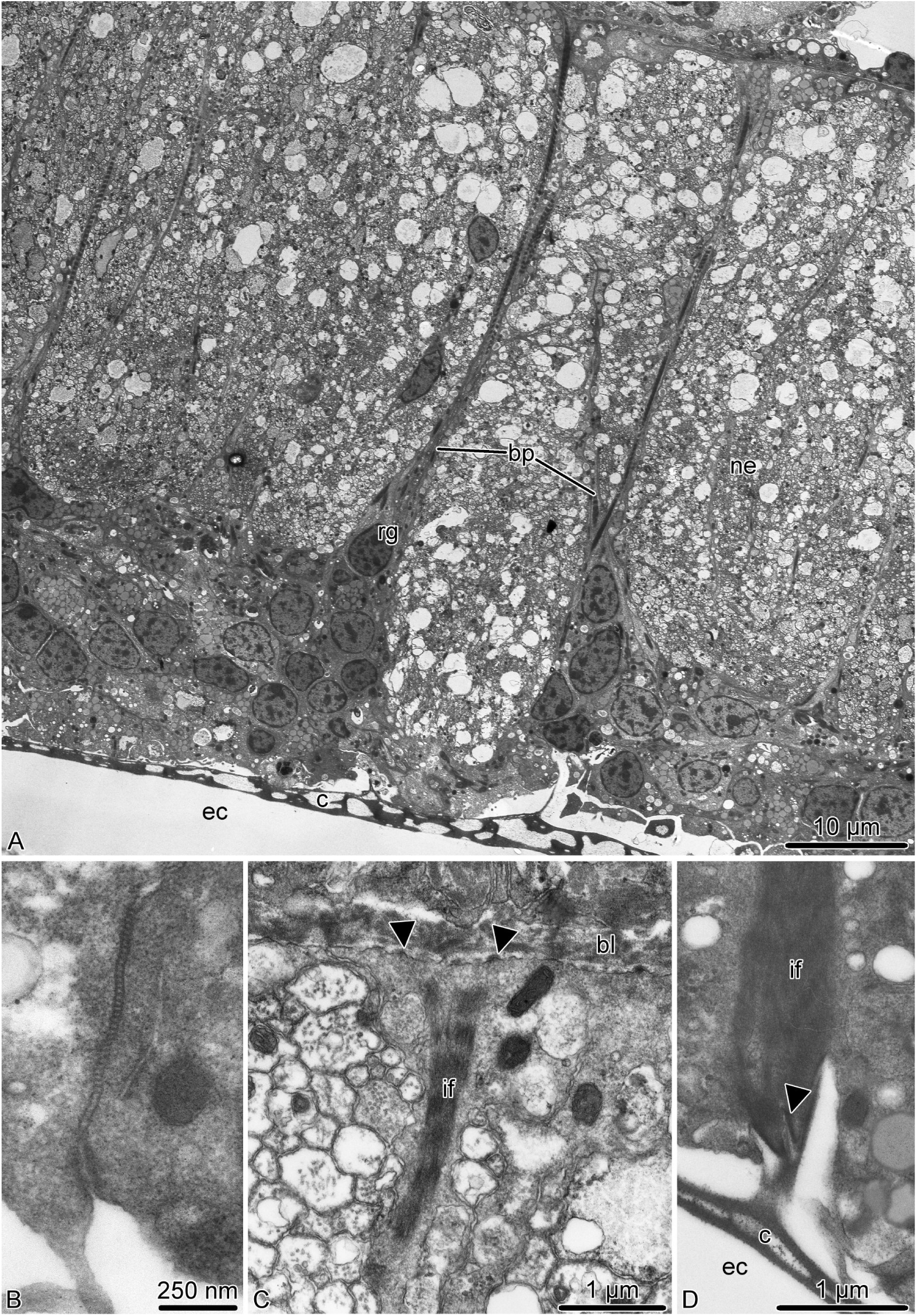
Radial glia in the ectoneural neuroepithelium of the radial nerve cord in *A. kochii*. Transmission electron microscopy. **A.** Low magnification view of the ectoneural neuroepithelium. **B.** Intercelluar junctions between apicolateral surfaces of two adjacent radial glial cells. **C.** Basal endfoot of a radial glial cell. **D.** Attachment of the apical surface of a radial glial cell to the epineural cuticle. Abbreviations: *bl* – basal lamina; *bp* – basal process of a radial glial cell; *c* – epineural cuticle; *ec* – epineural canal; *if* – intermediate filaments; *ne* – neuropil; *rg* – radial glia. *Arrowheads* show hemidesmosomes.

#### Ectoneural peripheral nerves and ganglia

At the level of the podia, the ectoneural part of the RNC gives off a pair of short and thick <u>podial nerves</u> (Fig. 3A, B, E; 4A, J). Shortly after emerging from the RNC, the podial nerve forms a thick ring ganglion at the base of the podium. A sleeve-like extension of the podial ganglion surrounds the hydrocoelic lining of the podium and descends up to the distal tip. The lateral side of the podial ganglion further gives off thick <u>spine nerves</u> to each spine (Fig. 3A, B; 4A, J). As they penetrate the lateral arm shields, these nerves pass through <u>spine ganglia</u>. The podial ganglia also give off small nerves that innervate tentacle scales (Fig. 4J).

#### Hyponeural peripheral nerves

The hyponeural part of the RNC gives off an extensive system of peripheral nerves (Fig. 3C, D). Although the hyponeural system by itself forms no purely hyponeural peripheral ganglia, it contributes to the formation of the mixed ganglia, known as juxtaligamental nodes (Wilkie, 1979) (see below). At the level of the ganglionic swellings of the RNC, immediately proximal to the podial nerves, a pair of large <u>proximal muscle nerves</u> ascend from the lateral regions of the hyponeural cord, enter the vertebral ossicle and arch towards the median surface of the aboral muscles attached to the proximal surface of the vertebra (Fig. 3D, F; 4B–D). There, it fuses with a wide ribbon-like <u>median hyponeural nerve</u> (Fig. 3D, F; 4D–G). This flat nerve runs between the median surface of the intervertebral muscles – both aboral and oral – and the intervertebral ligament and fuses with its oral margin with the hyponeural part of the RNC (Fig. 3D; 4H, I). It gives off numerous short branches on its surface that innervate both the intervertebral muscles and the mutable collagenous tissue of the intervertebral ligament in the central region of the arm (Fig. 3D, F; 4D–G).

On either side of the arm, a <u>horizontal intermuscular nerve</u> connects the median hyponeural nerves to the lateral juxtaligamental node (see below). This nerve runs between the intervertebral and lateral ligament in the space between the oral and aboral muscles (Fig. 3C, D; 4F, G).

Immediately behind the point of origin of the proximal muscle nerves, the hyponeural part of the RNC gives off two pairs of lateral nerves that both ascend aborally along the lateral surface of the oral intervertebral muscle (Fig. 3C, D; 4C–E, G). One of them – the <u>distal lateral nerve</u> – innervates the lateral arm shield, whereas the second one – the <u>proximal lateral nerve</u> – directly connects the radial nerve cord to the <u>lateral juxtaligamental node</u> (see below) (Fig. 4C–G).

#### Mixed (ecto-/hyponeural) peripheral nerves and ganglia

represent the third sub-division of the arm peripheral nervous system that cannot be classified as neither ectoneural nor hyponeural, as these structures are formed by contribution of both. All components of the mixed peripheral nervous system are embedded into the outer wall of the arm coelom. The most prominent structures of this mexed system are the paired large <u>lateral juxtaligamental nodes</u> that lie one on each side of the arm, midway between the oral and aboral surfaces (Fig. 3E; 4F, G). Each lateral node is formed by the fusion of three neural components: the lateral end of the horizontal intermuscular nerve, the aboral end of the proximal lateral nerve, and the distal margin of the podial ganglion (Fig. 3C–E; 4F, G). The first two structures originate from the hyponeural system, whereas the podial ganglion is ectoneural (Fig. 3A, B; 4A, D–G, J). The lateral juxtaligamental nodes give off numerous short branching protrusions that enter the adjacent collagenous tissue of the lateral ligament (Fig. 4F, G). They also give rise to two pairs of larger nerves, both thin and wide, which run in opposite directions in the lateral wall of the arm coelom (Fig. 3E, F; 4F, I). One pair – <u>the aboral mixed nerves</u> – ascend towards the aboral midline (Fig. 4F, I). The two <u>oral mixed nerves</u>, one on each side of the arm, descend towards the oral midline and fuse to form the <u>oral juxtaligamental node</u>, which innervates the oral ligament (Fig. 4H).

### 3.2 Cellular architecture of the nervous tissue

#### 3.2.1 Radial nerve cord

The ectoneural neuroepithelium of the RNC contains prominent radial glial cells (Fig. 5). The brittle star radial glia is very similar to their counterparts previously described in starfish and sea cucumbers (Viehweg et al., 1998; Mashanov et al., 2006, 2010, 2013). These cells are robustly labeled by the ERG1 antibody (Fig. 14A; 15E, F; 16B, C; 17B, C”’), which was raised against a sea cucumber glial antigen (Mashanov et al., 2010). They stretch throughout the height of the neuroepithelium between the apical and basal surfaces (Fig. 5A; 15E, F; 16C) and show clear epithelial cell features. Their cell bodies are often located at the apical surface of the neuroepithelium (Fig. 5A; 6A, B) and give off a long basal process that crosses the underlying neuropil (Fig. 5A; 6A, C). The distal end of the glial process forms a flattened endfoot that attaches to the basal lamina by hemidesmosomes (Fig. 5C). The cell bodies of adjacent glial cells are connected by apicolateral junctional complexes composed of zonula adherens and septate junctions (Fig. 5B). A typical intracellular characteristic of radial glia is the presence of thick bundles of intermediate filaments which run along the long axis of the cell (Fig. 5C, D). The epineural canal contains a dense accumulation of fibrous extracellular material that forms a flat cuticle-like structure overlaying the apical surface of the ectoneural neuroepithelium (Fig. 5A, D). This cuticle varies in thickness and sometimes shows a mesh-like organization of interconnected layers. The apical surface of the radial glial cells is attached to the cuticle via hemidesmosome-like contacts (Fig. 5D).

Most neuronal cell bodies are localized to the subapical region of the ectoneural neuroepithelium beneath the layer of glial cell bodies (Fig. 6A). Some neuronal perikarya, however, reach the lumen of the epineural canal (Fig. 6B). These neuronal cells are flanked by radial glia and joined to them by intercellular junctions. The region of the neuroepithelium located between the apical layer of the neuronal and glial cell bodies and the basal lamina is occupied by an extensive neuropil (Fig. 5A; 6C – F’). The neuropil is composed of densely packed neuronal processes filling all the space between the basal processes of radial glia (Fig. 6C). By their size, the neuronal processes can be clearly classified into ”regular” (measuring 130 nm – 600 nm across) and ”giant” (up to ∼5 *μ*m across) (Fig. 6C, D). Occasionally, the neuropil contains processes of neurosecretory-like cells (Fig. 6E) containing large dense granules, which are morphologically identical to those described in juxtaligamental cells (Wilkie, 1979). Well-defined chemical synapses are often seen in the neuropil (Fig. 6F, F’).

**Figure 6:**
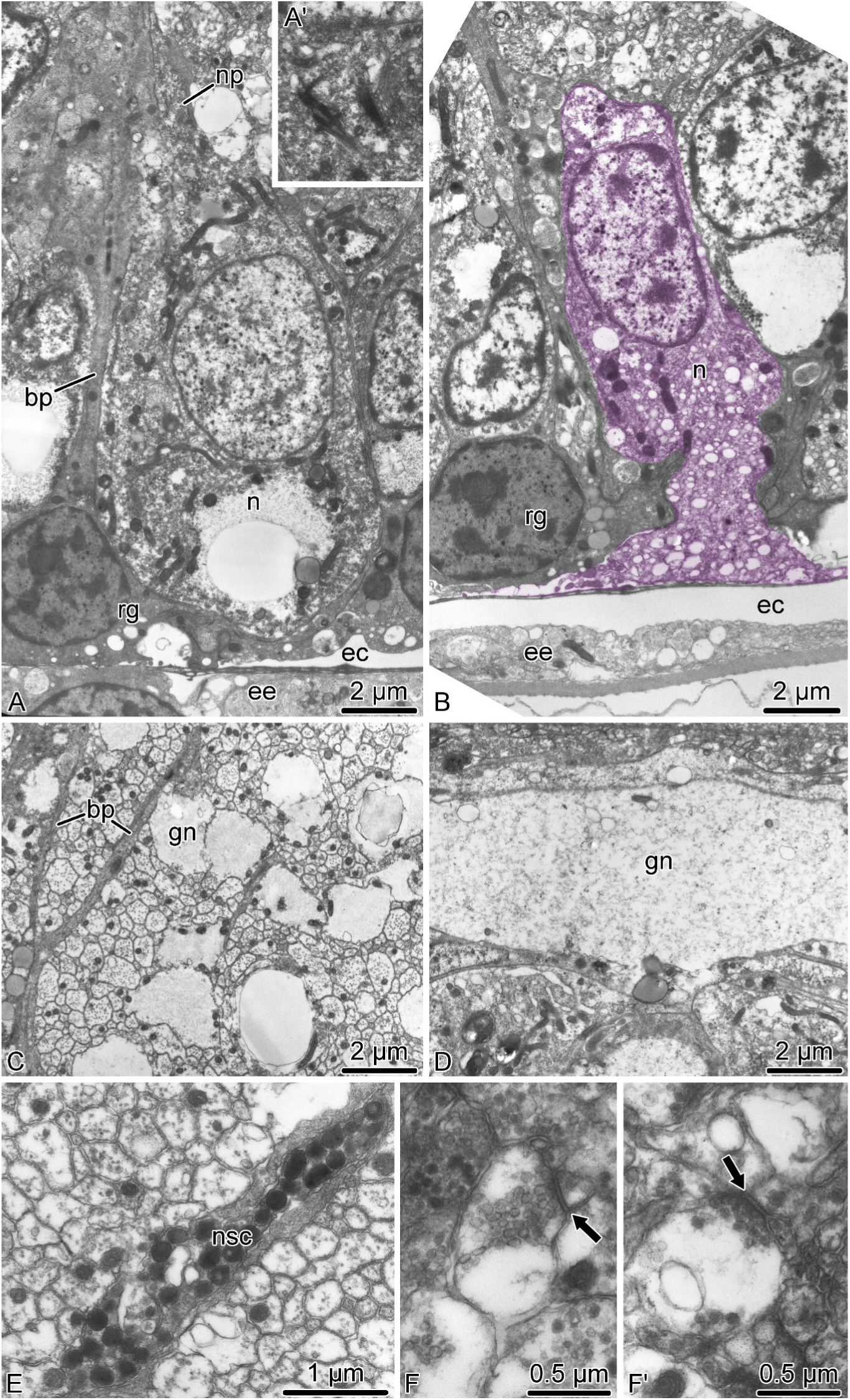
Neurons in the ectoneural neuroepithelium of the radial nerve cord in *A. kochii*. Transmission electron microscopy. **A.** Sub-apical neuron. A’. Cilium in a subapical neuron *(n)*. **B.** Neuron *(n, colored)* that reaches the lumen of the epineural canal *(ec)*. C – F’. Ectoneural neuropil. C and D show the transversal and longitudinal sections, respectively, of the neuropil area containing processes of giant neurons *(gn)*. **E.** Process of the neurosecretory-like cell *(nsc)*. F and F’. Synapses *(arrows)* in the neuropil. Abbreviations: *bp* – basal process of a radial glial cell; *ec* – epineural canal; *ee* – epineural epithelium (roof of the epineural canal); *gn* – giant neuronal processes; *n* – neuron; *np* – neural process; *nsc* – neurosecretory-like cell; *rg* – radial glial cell.

The roof of the epineural canal is formed by a thin epineural epithelium, a simple epithelial monolayer composed of flattened glial cells (Fig. 7A). Unlike the radial glia in the neuroepithelium, these glial cells mostly lack intermediate filaments, except those that are associated with the hemidesmosomes that anchor the cells to the basal lamina and to the cuticle in the epineural canal (Fig. 7B). Very occasionally, this epithelium contains basiepithelial nerve processes, but never neuronal perikarya (Fig. 7C).

**Figure 7:**
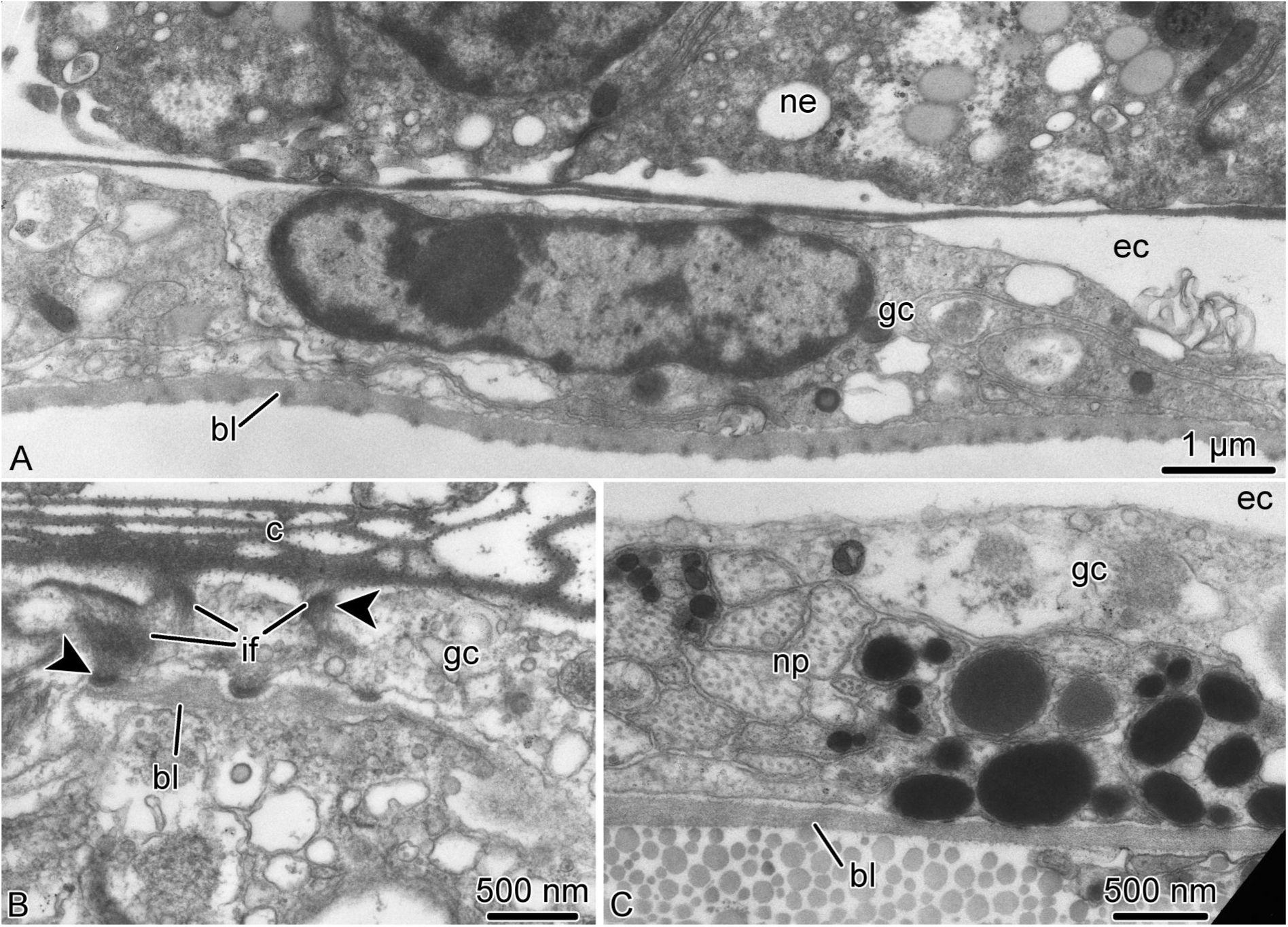
Roof of the epineural canal (epineural epithelium) in *A. kochii*. Transmission electron microscopy. **A.** Flattened glial cell *(gc)* of the epineural epithelium. **B.** Hemidesmosomes *(arrowheads)* anchoring a glial cell of the epineural epithelium to the basal lamina *(bl)* and to the epineural cuticle *(c)*. **C.** The epineural epithelium occasionally contains basiepithelial neural processes *(np)*, but not neuronal perikarya. Abbreviations: *bl* – basal lamina; *c* – epineural cuticle; *ec* – epineural canal; *gc* – flattened glial cell; *if* – intermediate filaments; *ne* – ectoneural neuroepithelium; *np* – neuronal processes.

The hyponeural component of the RNC overlays the aboral side of the ectoneural cord. Unlike in the ectoneural system, both the floor and the roof of the hyponeural canal are formed by neuroepithelia (Fig. 8). The tissue architecture of the hyponeural neuroepithelium markedly differs between the ganglionic swellings and the interganglionic regions of the RNC. In the interganglionic regions, neuronal elements are rare or entirely absents, and the wall of the hyponeural tube is composed of the flattened glial cells (Fig. 8A, B). In contrast, neuronal cell bodies and processes are abundant in the ganglionic swellings (Fig. 8C – F). The neuronal cell types are diverse and include ”regular” small neurons, ”giant” neurons (Fig. 8F), and neurosecretory-like cells (Fig. 8D). The bundles of nerve process mostly run parallel to the long axis of the RNC, but there are some that run transversally (Fig. 8E, F). The glial cells in the swollen ganglionic regions of the hyponeural cord have typical organization of radial glia, with the cell body positioned in the apical region of the neuroepithelium and a long basal process descending toward the basal lamina (Fig. 8C, D). However, unlike in the ectoneural part of the RNC, the hyponeural radial glia do not contain bundles of intermediate filaments in their cytoplasm. All glial cells in the hyponeural system are secretory. Their cytoplasm contains vacuoles filled with material of moderate electron density, which is released into the lumen of the hyponeural canal via exocytosis (Fig. 8B, C).

**Figure 8:**
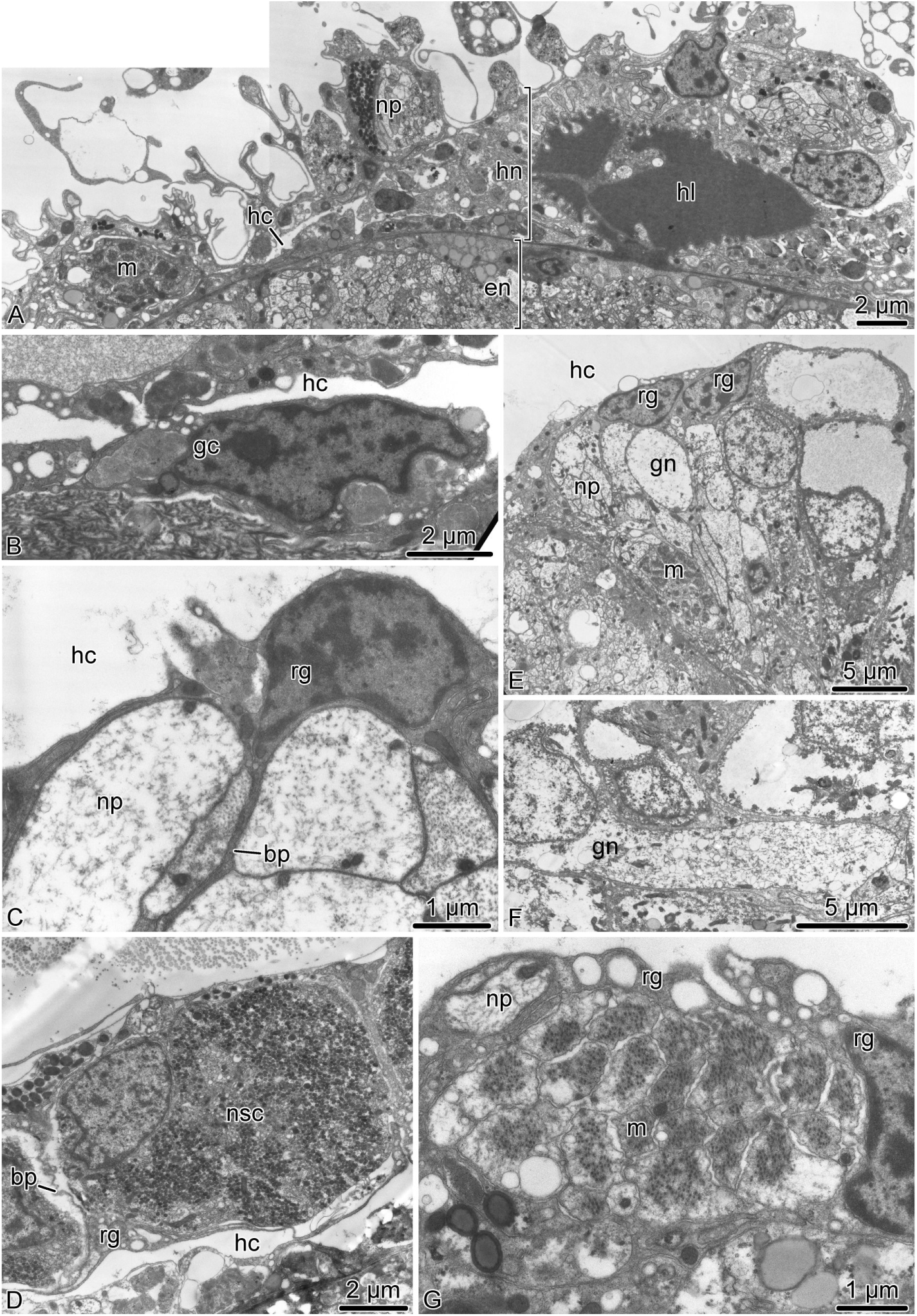
Organization of the hyponeural part of the radial nerve cord (RNC) in *A. kochii*. Transmission electron microscopy. **A** and **B.** Hyponeural system in the interganglionic region of the RNC. **A.** Low-magnification view of the hyponeural cord. **B.** Part of the oral wall (floor) of the interganglionic hyponeural cord formed by flattened glial cells with no neuronal elements. C – G Hyponeural system in the ganglionic swelling of the RNC. **C.** Secretory radial glia. **D.** Neurosecretory-like cell in the aboral wall (roof) of the hyponeural cord. E and F. Abundant neurons in the oral wall (floor) of the hyponeural cord. F. Giant neuron. G. High-magnification view of a muscle bundle integrated in the lateral region of the hyponeural neuroepithelium. Abbreviations: *bp* – basal process of radial glial cell; *en*– ectoneural neuroepithelium; *gc* – flattened glial cell; *gn* – ”giant” neuron; *hc* – lumen of the hyponeural canal; *hl* – hemal lacuna; *hn* – hyponeural neuroepithelium; *m* – bundle of muscle cells; *np* – neuronal processes; *nsc* – neurosecretory-like cell; *rg* – radial glia.

The hyponeural part of the brittle star RNC is also associated with two non-neural anatomical components. The first one is the radial hemal lacuna, which is a local expansion of the otherwise thin extracellular space that separates the ectoneural and hyponeural parts of the RNC (Fig. 8A). The lumen of the lacuna therefore does not have epithelial lining, but is instead surrounded by the basal lamina of the hyponeural neuroepithelium. The second non-neural component of the hyponeural system are two compact bundles of muscle cells immersed into the oral wall of the hyponeural cord on either side of the midline (Fig. 8A, E, G). These muscle bundles run longitudinally throughout the length of the radial nerve cord. As has been previously documented in studies of the sea cucumber CNS (Mashanov et al., 2006; Hoekstra et al., 2012), the ectoneural and hyponeural neuroepithelia of the RNC form extensive direct anatomical connections with each other. Even though the basal surfaces of these two neuroepithelia are separated from each other by a dense basal lamina, this separation is never complete. Frequent gaps in this basal lamina allow for the passage of neuronal processed from one neuroepithelium into another (Fig. 9A, A’). The basal lamina is also absent in the regions where the hyponeural part of the radial nerve cord comes in close contact with the oral intervertebral muscle (Fig. 9B, B’).

**Figure 9:**
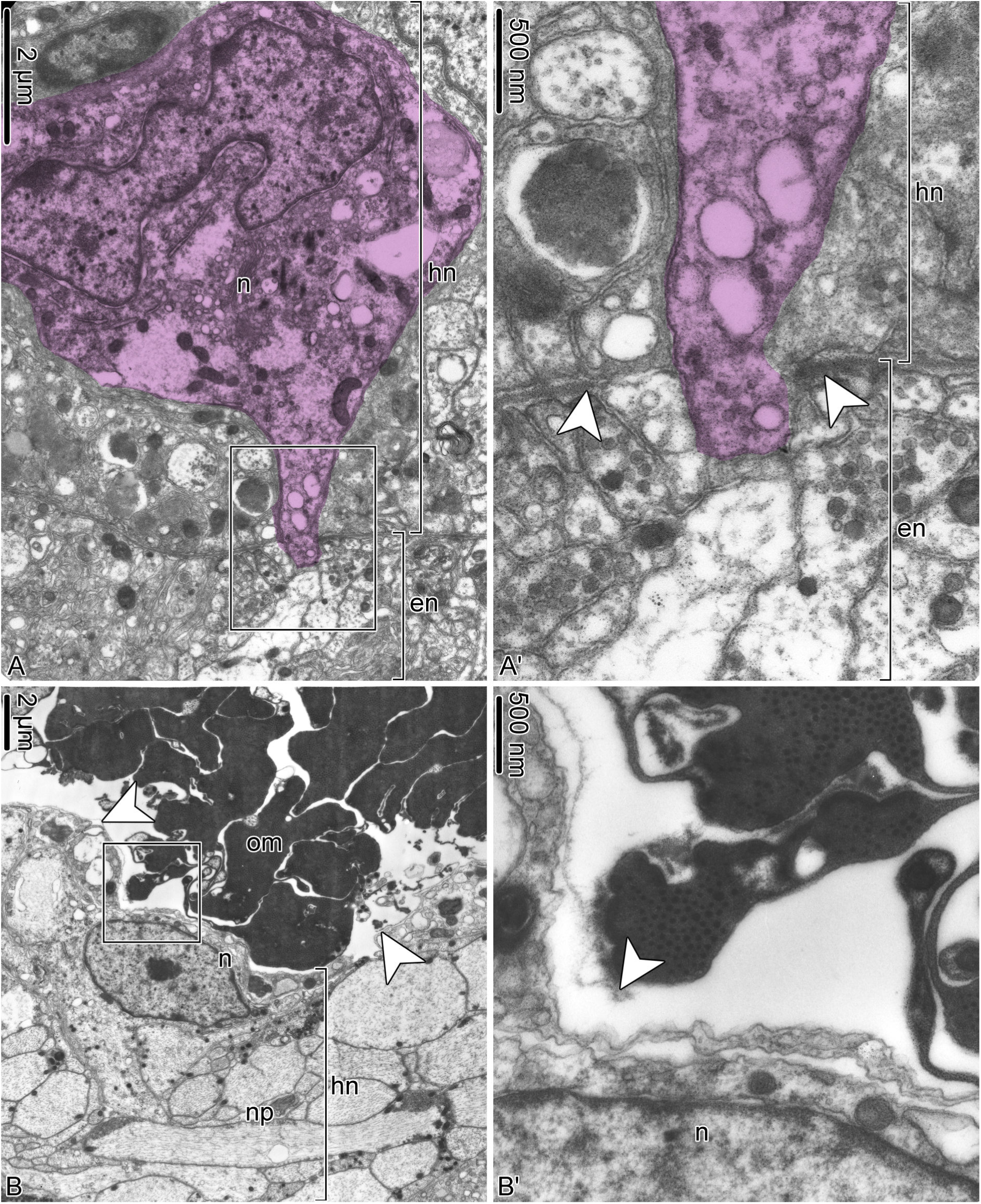
The hyponeural neuroepithelium makes direct contacts with the ectoneural epithelium (A and A’) and with the intervertebral muscles (B and B’) in *A. kochii*. Transmission electron microscopy. The colorized neuron in A and A’ has its cell body within the hyponeural neuroepithelium and sends a process into the ectoneural neuropithelium. A’ and B’ show detail views of the boxed areas in A and B, respectively. Abbreviations: *en* – ectoneural neuroepithelium; *hn* – hyponeural neuroepithelium; *n* – neuron; *np* – neural processes; *om* – oral intervertebral muscle. *White arrowheads* indicate the basal lamina.

#### 3.2.2 Peripheral nerves and ganglia

The ectoneural **spine ganglia** are formed by spherical clusters of bodies of neurosecretory-like juxtaligamental cells and neurons surrounding the ectoneural spine nerves (Fig. 4A, J; 10). There are two ultrastructurally distinct types of neurosecretory-like cells: those with abundant small spherical granules and those with larger less numerous ellipsoid granules (Fig. 10A). These neurosecretory cells are ciliated (Fig. 10D). The central neuropil of the spine ganglion is composed of the nerve processes of the spine nerve (Fig. 10B, C) and also contains well-defined chemical synapses between neuronal processes and neurosecretory cells (Fig. 10C). At the periphery, the spine ganglia are covered by a sheath of flattened glial cells and a basal lamina (Fig. 10E).

**Figure 10:**
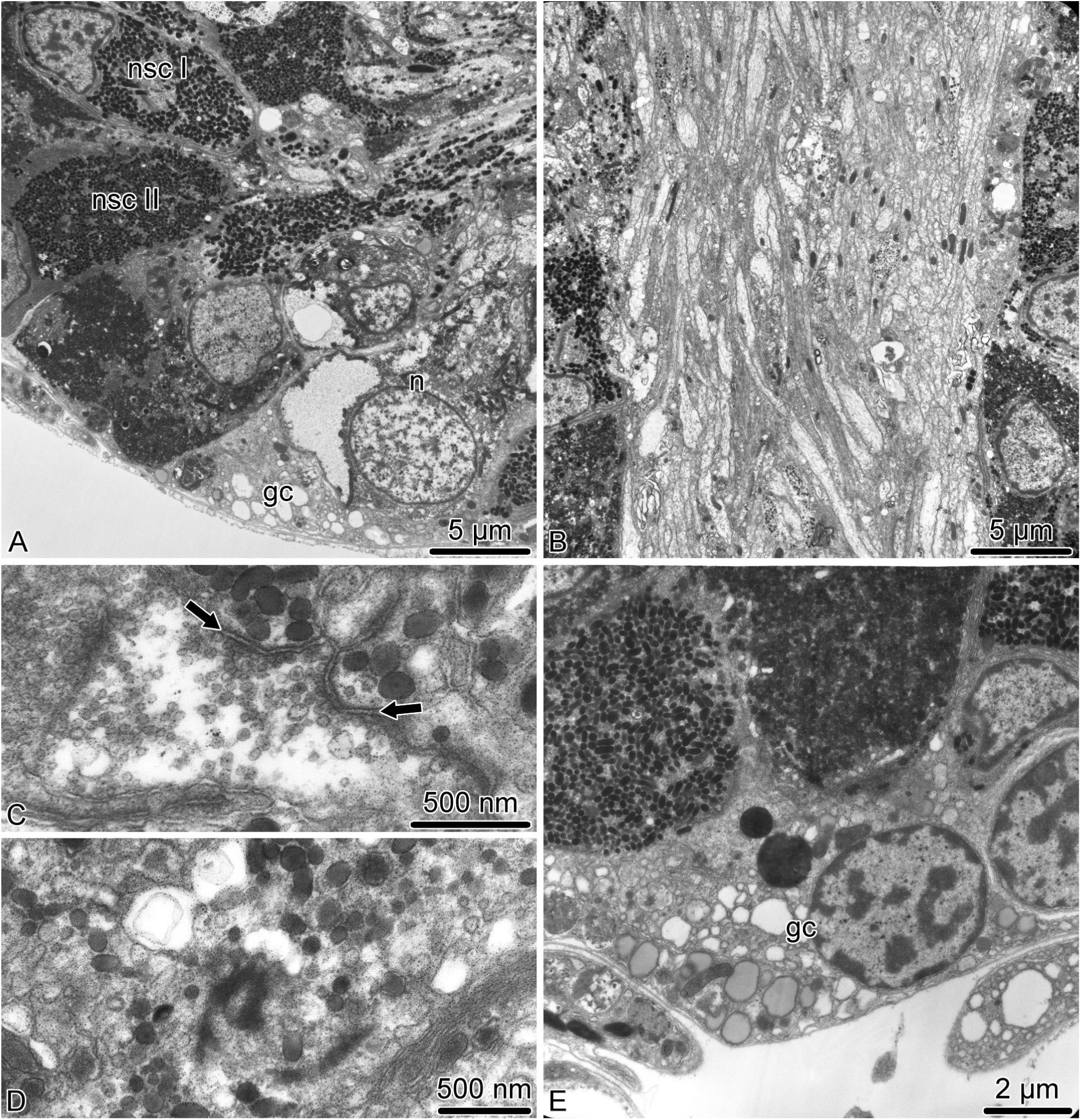
Organization of the spine ganglion in *A. kochii*. Transmission electron microscopy. **A.** Peripheral part of the spine ganglion with cell bodies of neurons and two types of neurosecretory-like cells. **B.** The central neuropil area of the spine ganglion with nerve processes of spine nerve passing through. **C.** Synapses *(arrows)* between a neural process and processes of neurosecretory-like cells. **D.** Basal body of a cilium in a neurosecretory cells. **E.** Glial cells at the periphery of the ganglion. Abbreviations: *gc* – glial cell; *n* – neuron; *nsc I* – neurosecretory-like cell type I; *nsc II* – neurosecretory-like cell type II.

The two major **hyponeural peripheral nerves** – the proximal muscle nerve and median nerve – have different ultrastructural organization. The proximal muscle nerve contains mostly thick processes of ”giant”neurons and occasionally processes of the ”regular” diameter and occasional neuronal perikarya (Fig 11A). The nerve is surrounded by a sheath of flattened glial cells and a continuous basal lamina. In contrast, the median hyponeural nerve lacks continuous glial envelope and has basal lamina only on the median side, which faces the intervertebral ligament (Fig 11B, C). On its lateral side, the nerve directly abuts the intervertebral muscle. The nerve processes are often seen to penetrate into the muscle and form synaptic contacts with myocytes (Fig 11B). The median nerves are particularly rich in cell bodies and processes of neurosecretory-like cells. The bundles of these processes frequently leave the nerve through the gaps in the basal lamina and branch in the collagenous connective tissue of the ligament (Fig 11C).

**Figure 11:**
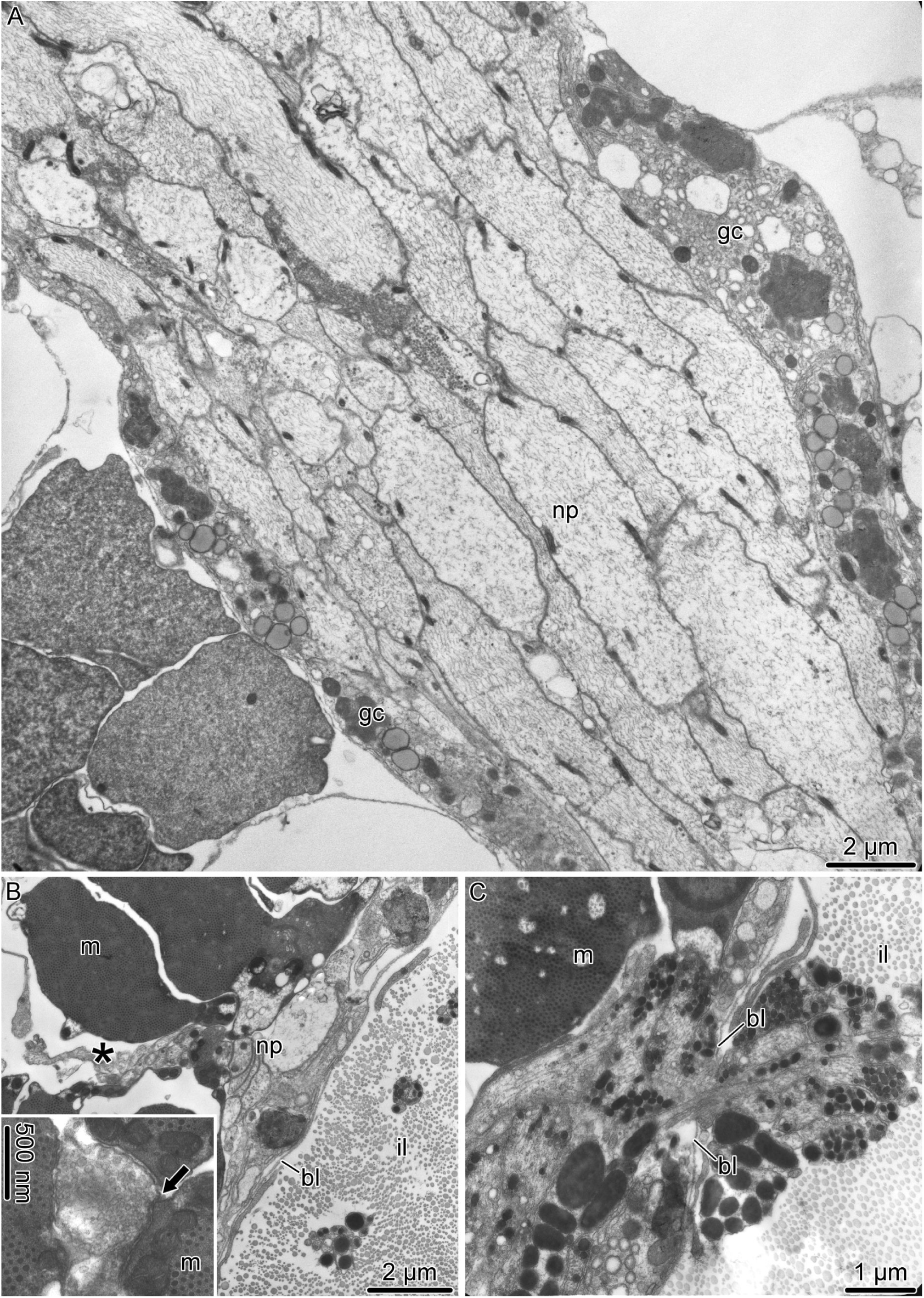
Organization of the hyponeural peripheral nerves in *A. kochii*. Transmission electron microscopy. **A.** Proximal muscle nerve. **B**, **C.** Median hyponeural nerve. **B.** Bundles of neuronal processes *(asterisk)* branching off the median hyponeural nerve and entering the oral intervertebral muscle. The inset in B shows a neuromuscular synapse *(arrow)*. **C.** Processes of neurosecretory-like cells passing through an opening in the basal lamina. Abbreviations: *bl* – basal lamina; *gc* – glial cell; *il* – intervertebral ligament; *m* – myocyte; *np* – neuronal processes.

Since all components of the **mixed peripheral nervous system** (i.e., the lateral and oral juxtaligamental nodes, aboral mixed nerves, and oral mixed nerves) are embedded into the outer wall of the arm coelom, they have typical organization of the basiepithelial nerve plexus. The oral and aboral mixed nerves are composed of cell bodies and processes of neurons and neurosecretory-like cells located between the cell bodies of the coelomic epithelial cells and the basal lamina of the coelomic epithelium (Fig. 12A, B). The ganglia – the oral and lateral juxtaligamental nodes – are local expansions of the basiepithelial plexus (Fig. 12C– H). They contain dense accumulations of cell bodies and extensive neuropil regions. The latter contain frequent synapses between neuronal processes and neurosecretory-like cells (Fig. 12E, H). Bundles of neurosecretory processes leave the ganglia through perforations in the basal lamina and enter the adjacent collagenous connective tissue (Fig. 12G).

**Figure 12:**
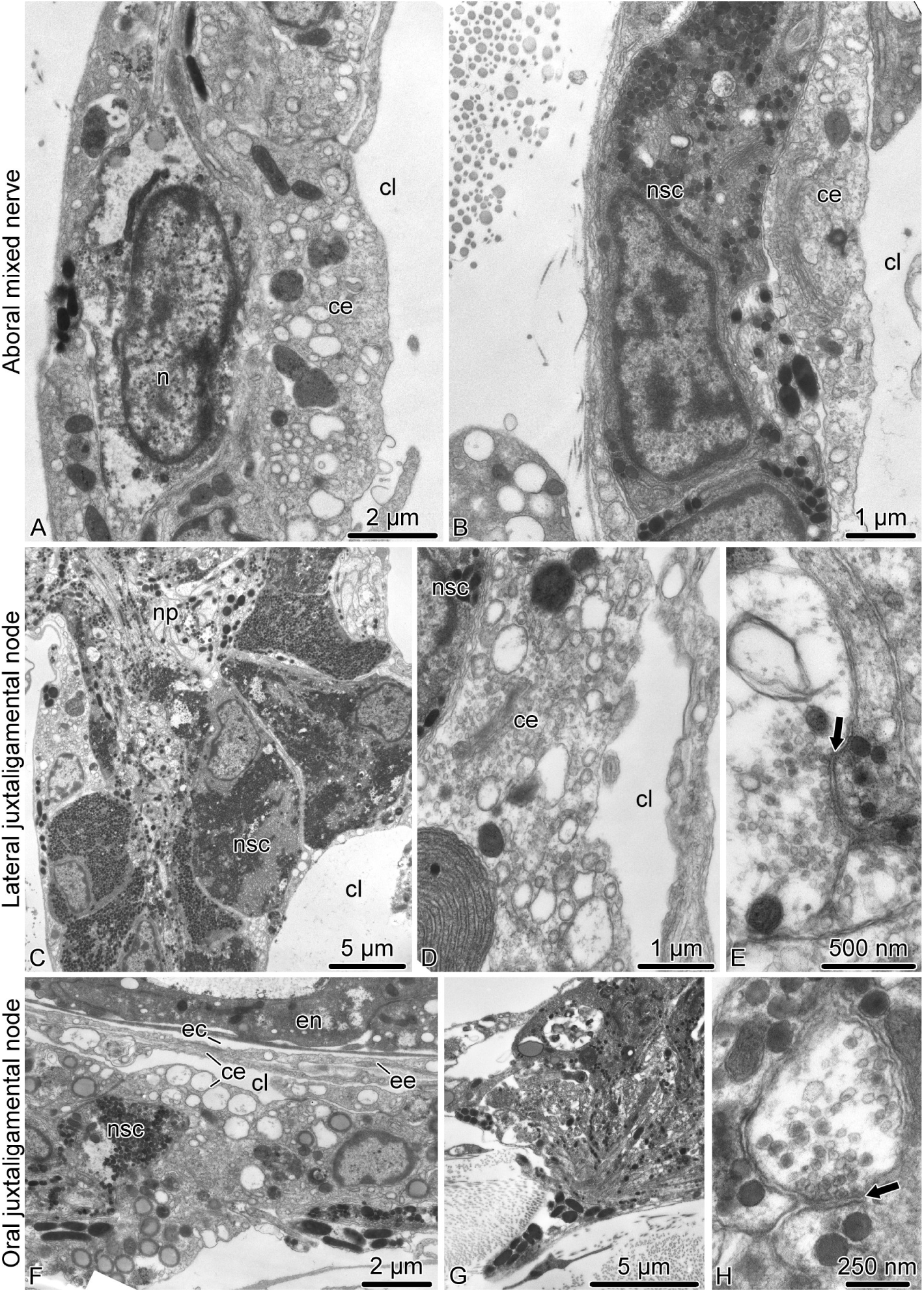
Organization of the mixed peripheral nerves and ganglia in *A. kochii*. Transmission electron microscopy. A, **B.** Aboral mixed nerve. C – **E.** Lateral juxtaligamental node. F – H. Oral juxtaligamental node. **A.** Neuronal cell body in the wall of the arm coelom.**B.** Neurosecretory-like cell in the wall of the arm coelom. **C.** Low magnification view of the lateral juxtaligamental node. **D.** Coelomic epithelial cell separating neurosecretory cells from the lumen of the coelom. **E.** Synapse *(arrow)* between an axon and a process of a neurosecretory cell in the neuropil area of the lateral juxtaligamental node. F. General view of the oral juxtaligamental node. G. Processes of neurosecretory-like cells leaving the oral juxtaligamental node and entering the collagenous connective tissue. H Synapse *(arrow)* between an axon and a process of a neurosecretory-like cell. Abbreviations: *ce* – coelomic epithelial cell; *cl* – lumen of the coelom; *ec* – epineural canal; *ee* – epineural epithelium; *en* – ectoneural neuroepithelium; *n* – neuron; *np* – neural processes; *nsc* – neurosecretory cell.

### 3.3 Immunohistochemistry

We used a number of cell-type specific markers to study localization of different cell types in the brittle star nervous system (Table 1). These include antibodies against general neuronal antigens, such as synaptotagmin B (SynB), ELAV, and acetylated tubulin; Brn1/2/4– an antigen specific to neuronal progenitors, and the neuropeptide GFSKLYFamide (Díaz-Miranda et al., 1995; Nakajima et al., 2004; Garner et al., 2016). To label glial cells, we used the ERG1 monoclonal antibody that stains radial glia in the echinoderm CNS (Mashanov et al., 2010).

The anti-SynB antibody extensively labels both the RNC and the peripheral nervous system of the arm (Fig. 13, 14). At the cellular level, it marks neuronal cell bodies that are mostly clustered in the proximal part of the ganglionic swollen regions of the RNC (Fig. 13A, A”, 14A, A’). In the RNC neuropil, it extensively stains neural processes, which mostly run longitudinally, in both the ectoneural and hyponeural systems. It also strongly labels the ring-shaped podial ganglia and a bundle of nerve processes that descends from the podial ganglion towards the tip of the podium on the median side (Fig. 13A, A’). The rest of the neural elements in the podium are stained as a fine net. Among other immunopositive structures in the peripheral nervous system are the spine nerves and cell bodies and fibers in the median hyponeural nerve that innervate the intervertebral muscles and ligaments (Fig. 13A, 14B–D). In the muscles, the SynB-containing processes form two parallel tracts, one near the proximal and one near the distal end. Individual fibers branch off those bundles and run parallel to the myocytes (Fig. 14B, C). The intervertebral ligament also contains a dense array of SynB-positive neural elements. These cells have small perikarya and give off long processes running parallel to the long axis of the ligament (Fig. 14D).

**Figure 13:**
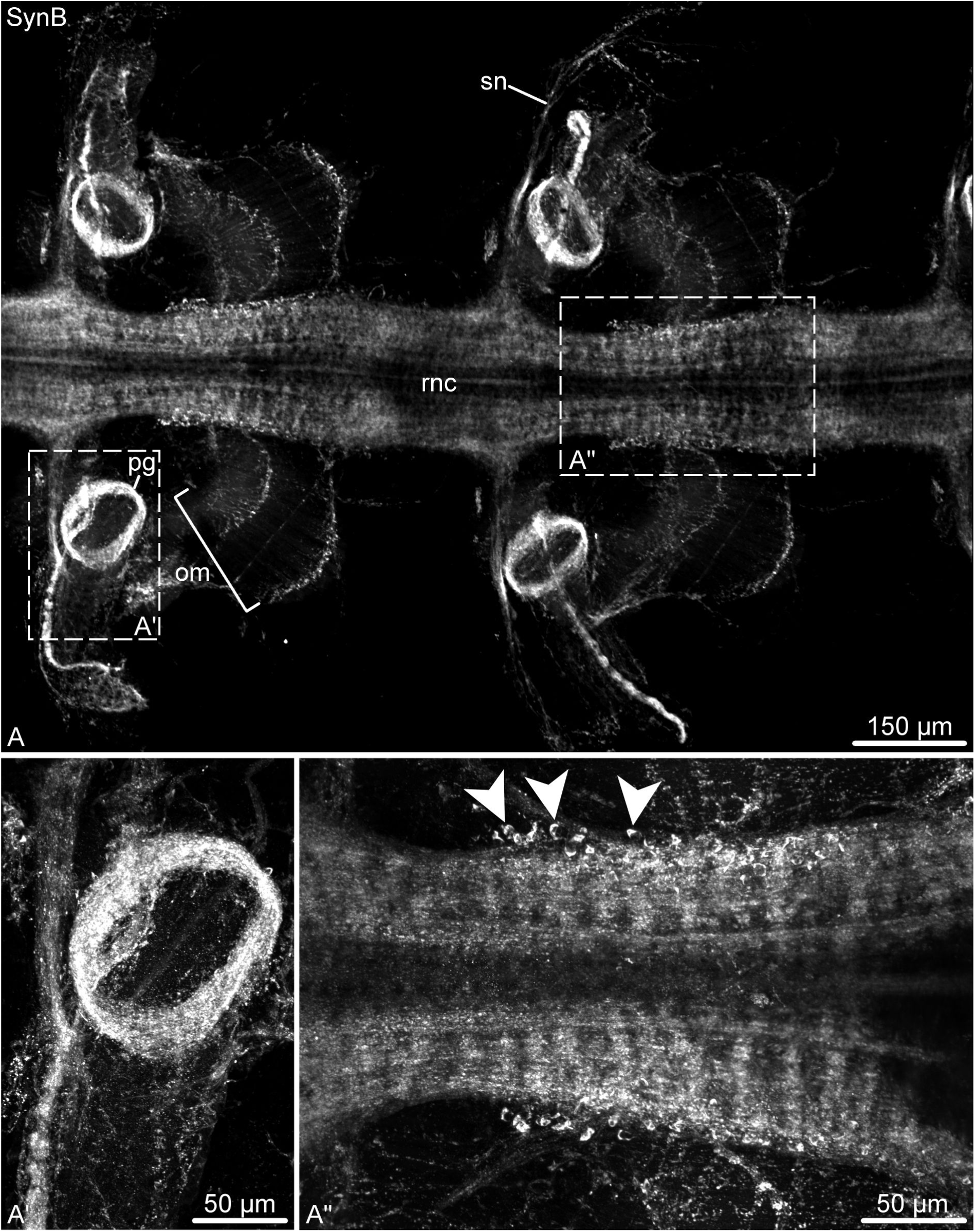
Synaptotagmin B (SynB) in the nervous system of the arm in *O. brevispinum*. Whole-mount preparation. Maximum intensity Z-projection of a confocal stack. A Low-magnification view of two segments of the radial nerve cord and associated peripheral nerves. A’ and A” show high-magnification views of the corresponding boxed areas in **A.** Abbreviations: *om* – oral intervertebral muscle; *pg* – podial ganglion; *rnc* – radial nerve cord; *sn* – spine nerve. *Arrowheads* in A” show neuronal cell bodies.

The anti-GFSKLYFamide antibodies label two well-defined longitudinal nerve tracts in the ectoneural neuropil of the RNC (Fig. 15A, A’, D, D’, F, F’). These tracts are positioned on either side of the midline and run continuously throughout the length of the arm. More loosely organized longitudinal processes also run on either side of the longitudinal tracts, but they do not form well-defined bundles (Fig. 15A, A’). In each ganglionic swelling of the RNC, the midline region between the two longitudinal tracts also contains a network of fibers (Fig. 15A–C), which largely dissipates in interganglionic regions. In each arm segment, the longitudinal GFSKLYFamide-positive tracts give off side branches that contribute to the podial nerves and podial ganglion (Fig. 15D, D’). The immunopositive neuronal cell bodies are not very numerous and are scattered in the ectoneural neuroepithelium without forming any noticeable clusters (Fig. 15A–C, E, E’). They lie at the apical region of the neuroepithelium. Some are clearly bipolar with an apical process reaching towards the lumen of the epineural canal and the basal axon descending in to the neuropil. Staining with the anti-GFSKLYFamide antibodies also reveals that at least some neurons in the RNC occupy stereotyped positions in different arm segments. For example, we identified a pair of large ectoneural unipolar neurons, which were always precisely localized to the distal region of the ganglionic swelling and projected their axons to the longitudinal tracts in the ectoneural neuropil (Fig. 15B, C).

**Figure 14:**
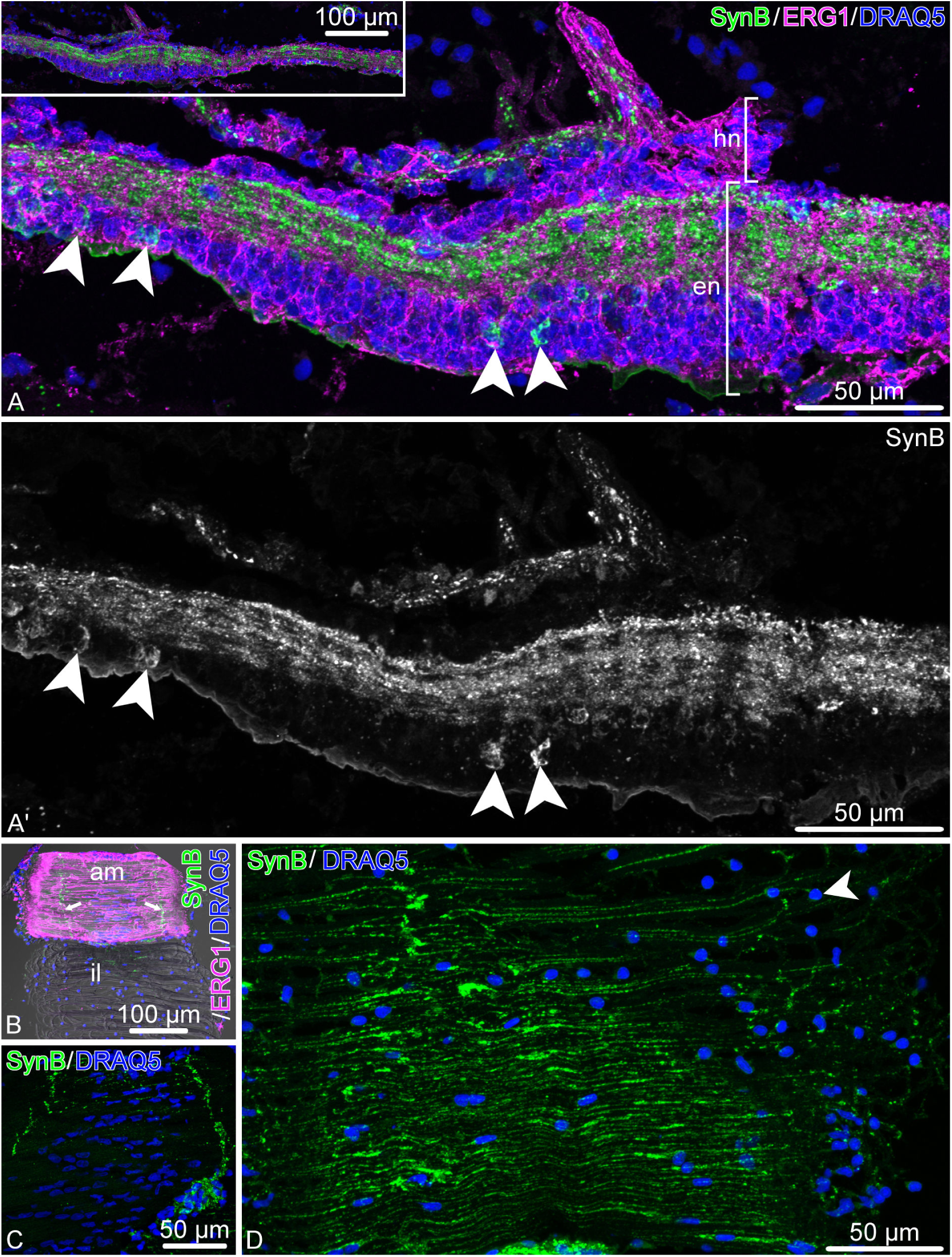
Synaptotagmin B (SynB) in the nervous system of the arm in *O. brevispinum*. Longitudinal sections. **A.** Radial nerve cord stained with the anti-SynB antibody and with the glial marker ERG1. The SynB staining is shown by itself in A’. The inset in A shows a low-magnification view of the radial nerve cord. **B.** Aboral intervertebral muscle and ligament. Note the immunopositive bundles of nerve fibers in the muscle *(arrows)*. **C.** Detailed view of innervation of the aboral muscle by SynB-immunopositive neurons. **D.** Innervation of the intervertebral ligament. Abbreviations: *am* – aboral muscle; *en* – ectoneural neuroepithelium; *hn* – hyponeural neuroepithelium; *il* – intervertebral ligament. *Arrowheads* in A, A’ and D how the cell bodies of SynB-immunopositive neurons.

**Figure 15:**
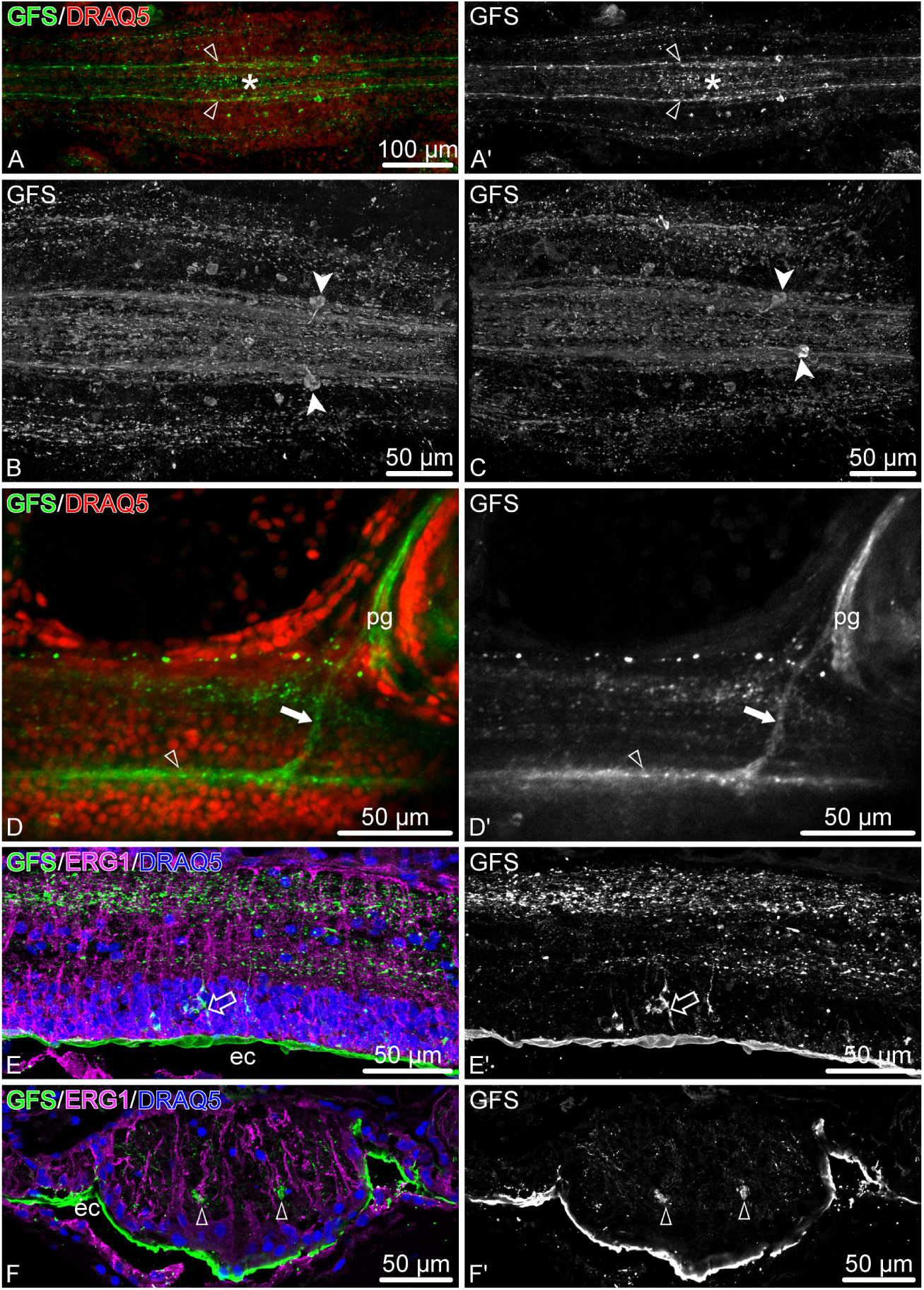
Distribution of the neuropeptide GFSKLYFamide (GFS) in the arm nervous system in *O. brevispinum*. **A and A’.** Low magnification view of a RNC segment. Whole-mount specimen, Z-projection of a confocal stack. **B and C.** A pair of stereotypically positioned neurons in two different arm segments. Whole-mount specimen, volume-rendered confocal stack. **D and D’** A branch of the longitudinal immunopositive tract contributing to the podial nerve. **E and E’.** Longitudinal section through the ectoneural neuroepithelium of the RNC. E shows triple staining with the anti-GFS antibodies, the ERG1 glial marker and the DRAQ5 nuclear stain, whereas E’ shows only the GFS-positive cells in a separate channel. **F and F’.** Cross section through the ectoneural epithelium of the RNC. F shows staining with the anti-GFS antibodies, ERG1 antibodies, and DRAQ5 nuclear dye, whereas F’ shows only anti-GFS immunostaining in a separate channel. Abbreviations: *ec* – epineural canal; *pg* – podial ganglion; *open arrowhead* – longitudinal immunopositive tracts; *asterisk* – median network of immunopositive fibers between the longitudinal tracts; *filled arrowheads* – a pair of stereotypically positioned neurons; *filled arrow* – a bundle of processes contributing to the podial nerve; *open arrow* – a bipolar neuron.

The neuron-specific RNA-binding protein ELAV is expressed in the majority of (or possibly all) neurons in both the ectoneural and hyponeural neuroepithelia of the RNC (Fig. 16). The anti-ELAV antibody also labels neurons in peripheral nerves, such as the podial nerve and the proximal muscle nerve (Fig. 16B, B’). The immunoreactivity appears to be specific to neurons, as ELAV-positive cells are never co-labeled by the radial glial marker ERG1 (Fig. 16D–D”’).

**Figure 16:**
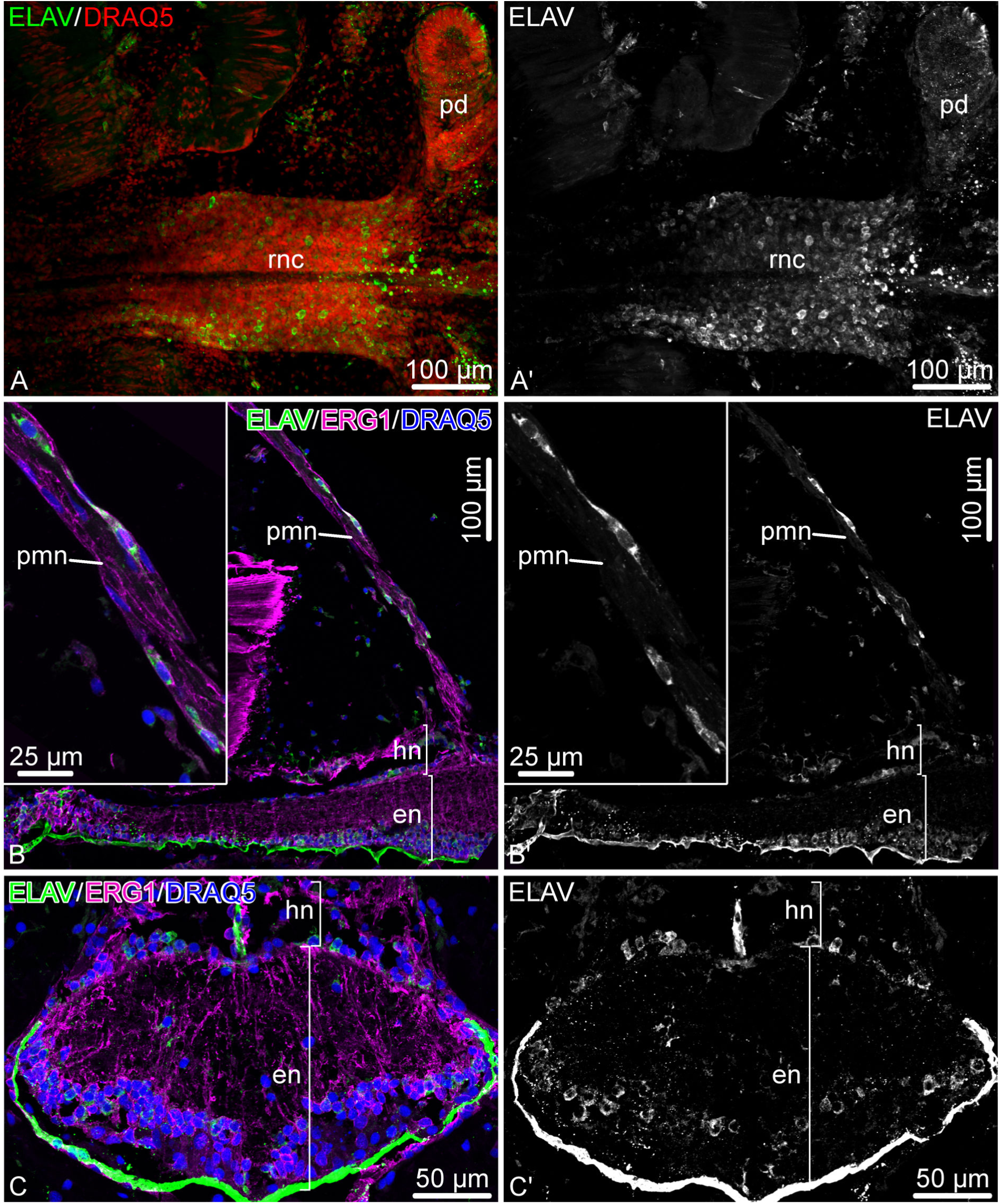
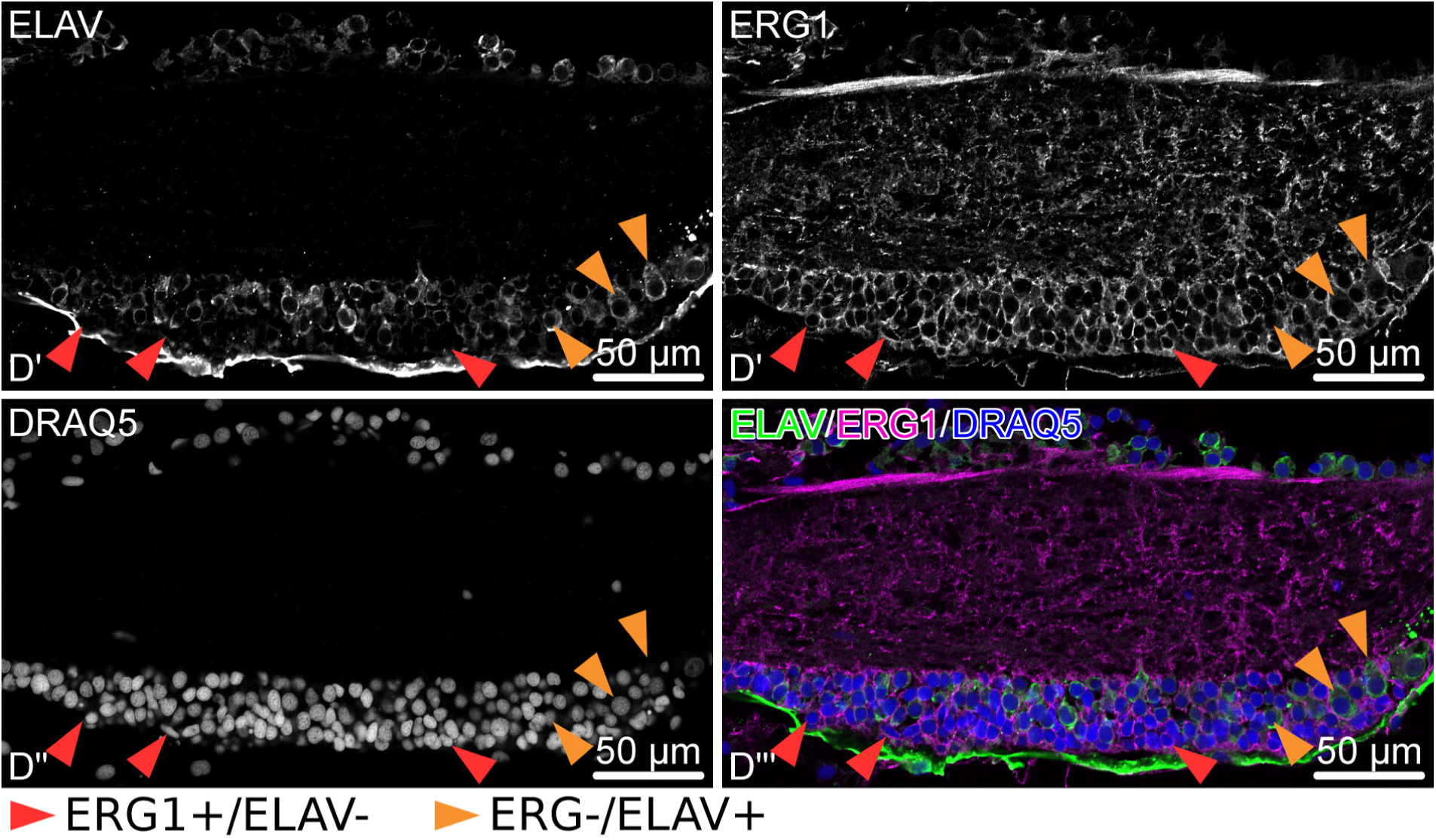
ELAV in the arm nervous system of *O. brevispinum*. **A and A’** Low magnification view of a RNC segment. Whole-mount specimen. Z-projection of a confocal stack. **B and B’.** Low-magnification view of a longitudinal section of the RNC. The insets show higher magnification of the proximal muscle hyponeural nerve. **C and C’.** Cross section of the RNC at the ganglionic swelling level. **D – D”’** High-magnification view of a longitudinal section of the RNC showing that there is no co-labeling of the same cells with the ERG1 and anti-ELAV antibodies. Abbreviations: *en* – ectoneural neuroepithelium; *hn* – hyponeural neuroepithelium; *pd* – podium; *pmn* – proximal muscle hyponeural nerve; *rnc* – radial nerve cord.

The transcription factor Brn1/2/4 is expressed in numerous cells of the ectoneural epithelium in the RNC. The expression is however restricted to cells of the ganglionic swelling (Fig. 17). Double immunostaining with the anti-Brn1/2/4/ antibody and the glial maker ERG1 shows that Brn1/2/4 is produced in both radial glial cells and neurons (Fig. 17C– C”’). Although all neurons (i.e., ERG1-negative cells) in the ganglionic swelling appear to express this transcription factor, there are two population of glial cells: Brn1/2/4-positive glia and Brn1/2/4-negative glia. In the ganglionic swelling these two types of glia are intermixed, whereas all glia in the interganglionic regions appear to be Brn1/2/4-negative.

**Figure 17:**
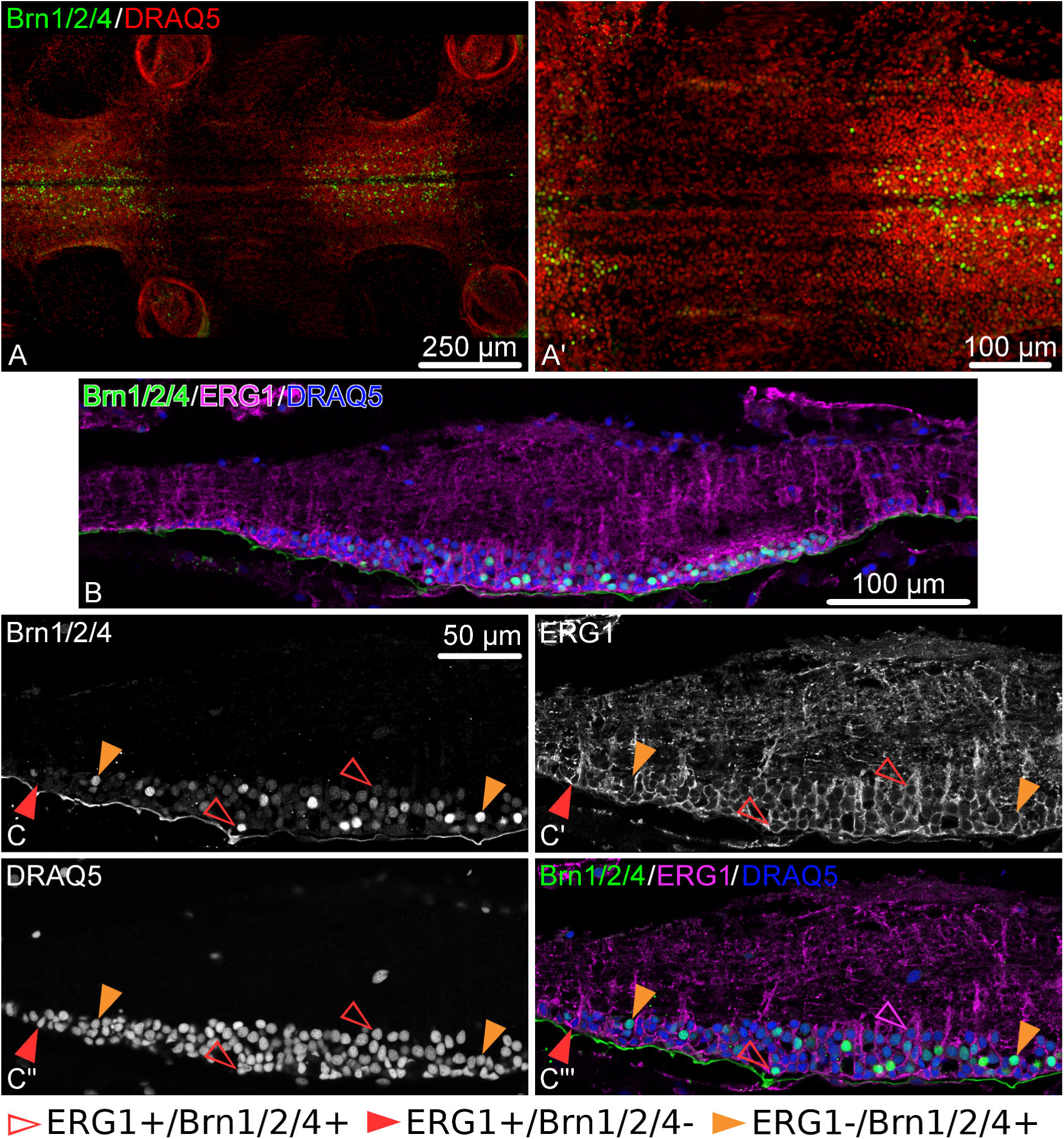
Expression of the transcription factor Brn1/2/4 in the radial nerve cord (RNC) of *O. brevispinum*. **A and A’** Whole-mount preparation showing the oral view of two RNC segments. Maximum intensity Z-projection of a confocal stack. **B.** A low-magnification view of a sagittal section through the RNC. **C–C”’.** High-magnification view of a longitudinal section of the RNC co-labeled with anti-Brn1/2/4 antibody and the ERG1 glial marker.

Acetylated tubulin is the marker that most robustly labels most of the components of the nervous system in the brittle star arm, both the RNC and peripheral nerves (Fig. 18, 19). In the RNC, it labels many – although not all – neuronal processes (Fig. 18). Many of the immunopositive neural fibers arise from the apically positioned neuronal perikarya. They descend into the neuropil and form mostly longitudinal bundles (Fig. 18C). Some fibers, however, are assembled into commissural tracts instead and cross into the contralateral side of the RNC (Fig. 18B). Even though the anti-acetylated tubulin antibody extensively labels neurites, it appears to stain only some, but not all perikarya. Many of the small neurons have an immunopositive axon, but no staining in the cell body. On the other had, ”giant”neurons have both their cell bodies and processes strongly labeled with this antibody (Fig. 18C).

**Figure 18:**
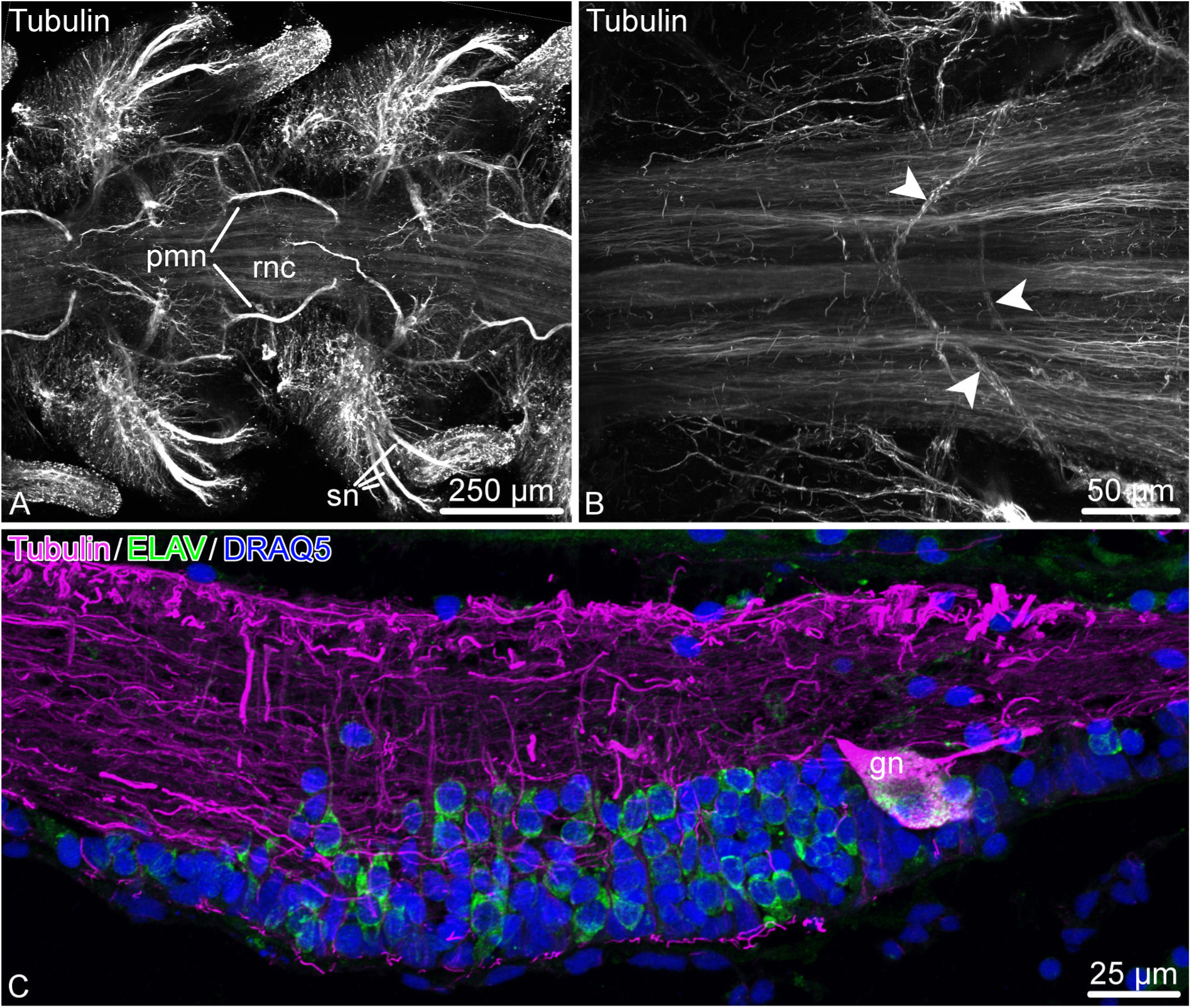
Acetylated tubulin in the radial nerve cord (RNC) of *O. brevispinum*. **A.** Low-magnification aboral view. Maximum intensity Z-projection of a confocal image stack. **B.** High-magnification view of neuronal fibers in the RNC. *Arrowheads* indicate commissural bundles crossing into the contralateral regions of the RNC. **C.** Sagittal section through the ganglionic swelling of the RNC. Dual labeling with anti-acetylated tubulin *(magenta)* and anti-ELAV *(green)* antibodies. The nuclei are stained with DRAQ. Note a ”giant” neuron *(gn)* in the distal region of the ganglionic region co-labeled with ELAV and acetylated tubulin. Abbreviations: *gn* – ”giant” neuron; *pmn* – proximal muscle nerve; *rnc* – radial nerve cord; *sn* – spine nerve.

**Figure 19:**
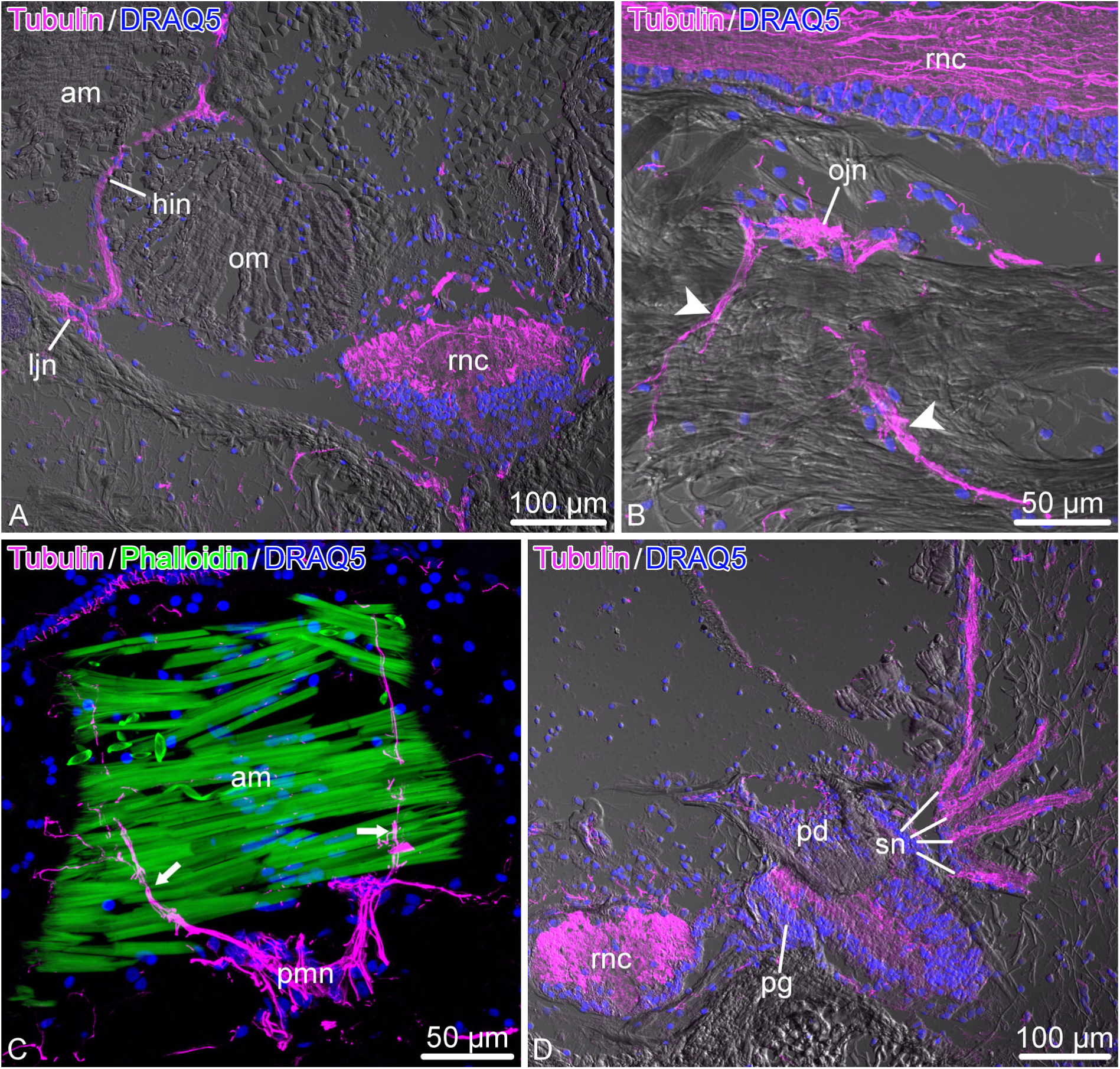
Acetylated tubulin in the peripheral nerves of the arm in *O.brevispinum*. **A.** Cross section showing the radial nerve cord, horizontal intermuscular hyponeural nerve, and lateral juxtaligamental node. **B.** Longitudinal section showing the oral juxtaligamental node and the bundles of immunopositive processes that originate from this ganglion and innervate the adjacent collagenous connective tissue *(arrowheads)*. **C.** Longitudinal section showing immunopositive neural processes *(arrows)* innervating the aboral muscle. **D.** Cross section through the podial ganglion and spine nerves. *sn*. Abbreviations: *am* – aboral muscle; *hin* – horizontal intermuscular hyponeural nerve; *ljn* – lateral juxtaligamental node; *ojn* – oral juxtaligamental node; *om* – oral muscle; *pd* – podium; *pg* – podial ganglion; *pmn* – proximal muscle nerve; *rnc* – radial nerve cord; *sn* – spine nerves.

Acetylated tubulin is also a reliable marker of the peripheral nervous system, as it marks both large nerves and their finest branches (Fig. 19). For example, it strongly stains the lateral and oral juxtaligamental nodes and the processes that these ganglia give off to innervate the adjacent regions of the collagenous connective tissue (Fig. 19A, B). It also marks the hyponeural proximal muscle nerve throughout its course, including the fibers that contribute to the intermusclular nerve and those that join the median hyponeural nerve to innervate the intervertebral muscles (Fig. 19C). Other peripheral nerves strongly marked with acetylated tubulin include the ectoneural spine nerves (Fig. 19D).

## 4 Discussion

In the recent years, there has been a renewed interest in revisiting the organization of the echinoderm CNS using modern techniques, including transmission electron microscopy and cell type-specific labeling to characterize individual glial and neuronal cell populations (Mashanov et al., 2006, 2010, 2016; Hoekstra et al., 2012; Díaz-Balzac et al., 2016). These recent studies resulted in a paradigm shift in our current understanding of echinoderm neurobiology and its phylogenetic significance. The echinoderm neural organization is no longer perceived as ”enigmatic”or unusual and is now considered to share a number of key features with other deuterostomes, including chordates. First, the ectoneural and hyponeural parts of the CNS, which were previously considered anatomically and functionally separate structures (Cobb, 1987, 1989, 1995), are now clearly established to be extensively crosslinked by direct neuronal connections (Mashanov et al., 2006; Hoekstra et al., 2012). Moreover, these two components of the nervous system originate from the same source in development (Mashanov et al., 2007). Second, the neurons were found to extensively communicate via ”classical” chemical synapses (Mashanov et al., 2006), which were previously considered to be absent in echinoderms. Another critical finding was that echinoderm CNS has neuroepithelial architecture with the scaffold composed of radial glial cells. These glial cells are similar to the chordate radial glia in a number of morphological and functional properties, including their function as neuronal progenitors in adult neurogenesis and neural generation (Viehweg et al., 1998; Mashanov et al., 2006, 2009, 2010, 2013, 2015a,b).

All these new significant findings, however, mostly emerged from studies of the sea cucumber CNS, one of the five existing classes in the phylum Echinodermata. It is therefore unclear whether or not these newly discovered principles are applicable to other echinoderm classes and whether they characterize the nervous system of the phylum in general. In this study, we establish that all general features mentioned above (radial glia scaffold in the neuroepithelium, frequent chemical synapses in the neuropil regions in the neuroepithelium, direct anatomical connections between the ectoneural and hyponeural systems) are also seen in the nervous system of brittle stars. However, there are some distinct features too that are present in ophuiroids, but not in the sea cucumber CNS. One such characteristic is the clearly defined segmental organization of the arm nervous system at the anatomical and cellular levels, as has been suggested in some earlier studies (Cobb and Stubbs, 1981; Bremaeker et al., 1997). At the anatomical level, the RNC is subdivided into ganglionic swellings separated by narrower interganglionic regions. In each segment, the peripheral nerves emerge from the same regions of the RNC and innervate the same effectors. At the cellular level, one can identify the stereotypically positioned individual cell bodies of giant neurons that contribute their axons to certain areas of the neuropil. This observation has several implications: (1) an existence of a patterning mechanism responsible for the segmentation in development/arm regeneration; (2) a degree of functional autonomy of the nervous system within each segment.

Even though the nervous system of brittle stars shows a number of distinct features that set them apart from other echinoderms, at the level of the class Ophiuroidea, the neuroanatomical design, including the arrangements of the peripheral nerves, appears to be remarkably stable and conserved, as it does not change across the three species studied so far, which represent three different ophiuroid families. Except for the minor differences, such as the number of spine nerves, the two species described in the present study – *O. brevispinum* and *A. kochii* – showed the same neuroanatomical architecture, which also matched the detailed descriptions by Hamman (1889) for *Ophioglypha albida*.

Another interesting feature of the brittle star nervous system is its intimate association with effectors, including the arm muscles and the collagenous connective tissue structures. The median hyponeural nerve is particularly remarkable in this regard, as it innervates both large intervertebral muscles and the adjacent intervertebral ligament. Not only it gives off numerous side branches penetrating the muscles, the nerve itself lacks glial covering and also is not separated from the muscle by a basal lamina. This intimate association between the nervous system and effectors probably provides the anatomical basis for the distinct locomotory behavior of brittle stars. Unlike in other echinoderms, tube feet play a relatively minor role in the whole-body movement of brittle starts. Instead, they extensively use bending of their jointed arms to crawl over the substratum (Astley, 2012). These movements can be unusually rapid (by echinoderm standards) and are highly coordinated across the individual arms.

Direct motor control of the collagenous connective tissue is a unique echinoderm phenomenon (Wilkie, 2005). Neurosecretory-like cells, which in brittle stars are called juxtaligamental cells, have been consistently found in association with connective tissue structures capable of changing their mechanical properties. The reversible change has been reported to be involved in movement and posture control, while irreversible loss of tensile strength in ligaments and tendons is the main mechanism of CNS-controlled autotomy in echinoderms. The changes in the properties of the extracellular matrix are mediated by substances released by juxtaligamental cells. These neurosecretory cells, in turn, are believed to be directly innervated by the central nervous system. Here, we provide direct support to this model, as we consistently see direct chemical synapses formed by neuronal terminals on juxtaligamental cells. Previously, the cell bodies of juxtaligamental cells were mainly found in peripheral ganglia (juxtaligamental nodes) in the direct vicinity of the mutable collagenous structures they control (Wilkie, 1979, 2005). Here, we found cells of identical ultrastructural appearance in the RNC. This suggests that there are two cohorts of juxtaligamental cells: peripheral juxtaligamental cells localized in the vicinity of the structures they control and the central juxtaligamental cells with the cell bodies localized in the RNC and processes contributed to peripheral nerves. The respective physiological roles of these two populations of neurosecretory cells remains to be established.

A pair of small bundles of muscles cells are also incorporated into the oral wall of the hyponeural part of the RNC and run throughout the length of the nerve cord. These bundles of myocytes within the RNC are unique features of the brittle star nervous system, as they have never been observed in other echinoderms. The purpose of these cells is not known, but their position and organization allows us to formulate some preliminary thoughts. These bundles are very small, especially in comparison with the powerful intervertebral muscles. The cytoskeletal components of their contractile apparatus are weakly developed. Finally, unlike other muscles in the arm, these myocytes are never connected to any of the skeletal elements in the arm. Instead, they are completely immersed into the nervous tissue and fully surrounded by glial and neuronal cells. Taken together, these three observations suggest that these bundles of myocytes are highly unlikely to generate any significant contractile force. We hypothesize that they instead may function as stretch receptors (proprioceptors) immersed into the CNS. It would be interesting to experimentally probe into the function of these cells and to trace their origin in development and regeneration.

Another important finding that emerged from this study is that it contributes evidence in support of the idea that the glia in echinoderms are diverse and heterogeneous. Previous studies of the sea cucumber CNS demonstrated that the radial glial cells in the RNC, in spite of being all morphologically alike, fell into two distinct subpopulations: some of the glia expressed the transcription factor Myc, while in others it remained transcriptionally silent (Mashanov et al., 2015a). Here, we show that in the brittle star RNC, radial glial cells also differ in their expression of the Brn1/2/4 transcription factor and thus form distinct Brn1/2/4^+^ and Brn1/2/4^-^ subpopulations. These two glial subtypes are intermixed within the ganglionic swellings of the RNC, but the interganglionic regions contain only the Brn1/2/4^-^ glia. Besides glia, Brn1/2/4 is also expressed in all neurons within the ganglionic swellings and thus its expression is thus not turned off in fully mature brittle star neurons. Brn proteins are a subgroup of the POU family transcription factors. In vertebrates, they have been implicated in neurogenesis and specification of the neuronal fate (Vierbuchen et al., 2010; Brombin et al., 2011). The neurogenic function appears to be evolutionary conserved, as Brn1/2/4 was expressed in post-mitotic differentiating neuronal progenitors in the developing larval nervous system of a sea urchin (Garner et al., 2016). The functional significance of Brn1/2/4 in a subset of glial cells remains unclear. One possibility is that this transcription factor marks the neurogenic population of radial glia. It has been previously shown that radial glia in sea cucumbers give rise to both neurons and new glial cells in neural regeneration, but also in the uninjured adult CNS (Mashanov et al., 2013, 2015a). Even though neurogenesis is negligible in the brittle star RNC, radial glia undergoes rapid activation followed by extensive cell proliferation after arm autotomy (Mashanov et al., in preparation). It therefore remains to be established if the differences in gene expression is related to the potency of radial glial cells in post-traumatic neurogenesis.

An additional level of echinoderm glial complexity involves the fact that glial cells are not restricted to the CNS only. Here, we confirm the existence of the peripheral glial cells, previously reported by Byrne (1994), which are associated with some peripheral nerves (e.g. the hyponeural proximal nerve) and ganglia (e.g. spine ganglia).

Several brittle star species (Astley, 2012; Czarkwiani et al., 2016; Long et al., 2016; Delroisse et al., 2017; Zandawala et al., 2017) are currently being developed as model organisms to address various biological questions, ranging from evolutionary biology to developmental and behavioral biology. Solid understanding of ophiuroid neurobiology is often required for the success of those various projects. In turn, the availability of the cell-type specific markers is critical for studies of the organization and development of the nervous system. We, therefore, tested the suitability of a number of available antibodies (Table 1), both commercial and received as a gift from collaborators, for identification of specific cell populations in the central and peripheral nervous system in a brittle star. Acetylated tubulin, ELAV, and synaptotagmin B appear to be the best neuronal markers, whereas Brn1/2/4 also labels some, although not all, glial cells, as indicated above. The echinoderm glial marker ERG1 (Mashanov et al., 2010) reliably marks radial glia in the *O. brevispinum* and also the peripheral glia associated with, e.g., the proximal muscle nerve.

## 5 Conclusions

- Our results in combination with already available data on the sea cucumber nervous system enhance our understanding of general principles of echinoderm nervous system organization including:
  – the ectoneural and hyponeural components of the nervous system are extensively interconnected
  – the CNS has neuroepithelial organization with the supporting scaffold formed by radial glial cells
  – radial glial cells in the CNS are molecularly and probably functionally diverse
- The brittle star CNS is highly metameric. The same pattern of peripheral nerves/ganglia and precisely positioned bodies of at least some cells are repeated in all arm segments.
- For the first time, we have described a system of putative proprioceptors that are associated with the CNS and embedded into the hyponeural neuroepithelium.
- We tested the suitability of glial and neuronal markers for studies of the brittle star CNS. As expected, the radial glial cells reliably marked with the ERG1 antibody, whereas the best neuronal markers are acetylated tubulin, ELAV, and synaptotagmin B. The transcription factor Brn1/2/4, a marker of neuronal progenitors, is expressed in all neurons of the ganglionic swellings of the RNC, but also in a subset of glial cells.

## 6 Acknowledgments

The authors thank Dr. Robert Burke (University of Victoria, Canada) and Dr. José Garcίa-Arrarás (University of Puerto Rico) for their gift of the antibodies. We are also grateful to Ms. Beate Aschauer (LMU, Munich) for her technical assistance and to all members of the Mashanov lab at UNF for critical discussion and inspiring comments. The study was supported by the Alexander von Humboldt Foundation and University of North Florida.

## 9 Supplementary Files

### Supplementary File 1

Aligned stack (saved as a video *.avi* file) that was used to generate the 3D model. The movie progresses in the proximal-to-distal direction.

### Supplementary File 2

The 3D model of the arm nervous system (one segment is shown only) saved as a *.blend* file. The file can be opened and manipulated in Blender, a free open-source 3D editor (https://www.blender.org/). The following anatomical structures are represented: *green* – ectoneural system; *magenta* – hyponeural system; *light blue* – mixed peripheral nerves; *brown* – intervertebral muscles; *red* – water-vascular system (hydrocoel). A side of the scale cube measures 100 *μ*m.

### Supplementary File 3

Ectoneural system. 3D animation.

### Supplementary File 4

Hyponeural system. 3D animation.

### Supplementary File 5

All components of the arm nervous system. 3D animation.

### Supplementary File 6

All components of the arm nervous system (*green* – ectoneural system; *magenta* – hyponeural system; *light blue* – mixed peripheral nerves) plus the intervertebral muscles (*brown*) and hydrocoel (*red*). This is an interactive 3D model generated from the original *.blend* file (Supplementary File 2) using the Blend4Web tool (https://www.blend4web.com). The model can be opened in any modern web browser. Upon opening, zoom out using the scroll wheel. To rotate, press and hold the left mouse button. To pan the view, press and hold the right mouse button.

### Supplementary File 7

Ectoneural system. Interactive 3D model. See caption to Supplementary File 6 for instructions.

### Supplementary File 8

Hyponeural system. Interactive 3D model. See caption to Supplementary File 6 for instructions.

### Supplementary File 9

Mixed nerves. Interactive 3D model. See caption to Supplementary File 6 for instructions.

### Supplementary File 10

Arm hydrocoel. Interactive 3D model. See caption to Supplementary File 6 for instructions.

### Supplementary File 11

Intervertebral muscles of the arm. Interactive 3D model. See caption to Supplementary File 6 for instructions.

## References

Henry C. Astley. Getting around when you’re round: quantitative analysis of the locomotion of the blunt-spined brittle star, ophiocoma echinata. Journal of Experimental Biology, 215(11):1923–1929, 2012. ISSN 0022-0949. doi:10.1242/jeb.068460. URL http://jeb.biologists.org/content/215/11/1923.

Richard Bannister, Imelda M. McGonnell, Anthony Graham, Michael C. Thorndyke, and Philip W. Beesley. Coelomic expression of a novel bone morphogenetic protein in regenerating arms of the brittle star amphiura filiformis. Development Genes and Evolution, 218(1):33, Dec 2007. ISSN 1432-041X. doi:10.1007/s00427-007-0193-9. URL https://doi.org/10.1007/s00427-007-0193-9.

Nancy D. Bremaeker, Dimitri Deheyn, Michael C. Thorndyke, Fernand Baguet, and Jerome Mallefet. Localization of s1– and s2–like immunoreactivity in the nervous system of the brittle star amphipholis squamata (delle chiaje 1828). Proceedings of the Royal Society of London B: Biological Sciences, 264(1382):667–674, 1997. ISSN 0962-8452. doi:10.1098/ rspb.1997.0095. URL http://rspb.royalsocietypublishing.org/content/264/1382/667.

Alessandro Brombin, Jean-Philippe Grossier, Aurlie Heuz, Zlatko Radev, Franck Bourrat, Jean-Stphane Joly, and Franoise Jamen. Genome-wide analysis of the pou genes in medaka, focusing on expression in the optic tectum. Developmental Dynamics, 240(10):2354–2363, 2011. ISSN 1097-0177. doi:10.1002/dvdy.22727. URL http://dx.doi.org/10.1002/dvdy.22727.

M Byrne. Microscopic Anatomy of Invertebrates, chapter Ophiuroidea, pages 247–343. Wiley-Liss: New York, 1994.

J. L. S. Cobb. Neurobiology of the Echinodermata, pages 483–525. Springer US, Boston, MA, 1987. ISBN 978-1-4613-1955-9. doi:10.1007/978-1-4613-1955-9 17. URL https://doi.org/10.1007/978-1-4613-1955-9_17.

J. L. S. Cobb. The nervous systems of Echinodermata: Recent results and new approaches, pages 407–424. Birkhaüser Basel, Basel, 1995. ISBN 978-3-0348-9219-3. doi:10.1007/ 978-3-0348-9219-3 18. URL https://doi.org/10.1007/978-3-0348-9219-3_18.

J. L. S. Cobb and T. R. Stubbs. The giant neurone system in ophiuroids. Cell and Tissue Research, 219(1):197–207, Aug 1981. ISSN 1432-0878. doi:10.1007/BF00210028. URL https://doi.org/10.1007/BF00210028.

James L. S. Cobb. Enigmas of Echinoderm Nervous Systems, pages 329–337. Springer US, Boston, MA, 1989. ISBN 978-1-4899-0921-3. doi:10.1007/978-1-4899-0921-3 23. URL https://doi.org/10.1007/978-1-4899-0921-3_23.

Anna Czarkwiani, Cinzia Ferrario, David Viktor Dylus, Michela Sugni, and Paola Oliveri. Skeletal regeneration in the brittle star amphiura filiformis. Front Zool, 13:18, 2016. doi:10.1186/s12983-016-0149-x. URL http://dx.doi.org/10.1186/s12983-016-0149-x.

Jérome Delroisse, Esther Ullrich-Lüter, Stefanie Blaue, Olga Ortega-Martinez, Igor Eeckhaut, Patrick Flammang, and Jérome Mallefet. A puzzling homology: a brittle star using a putative cnidarian-type luciferase for bioluminescence. Open Biology, 7(4), 2017. doi:10.1098/rsob.160300. URL http://rsob.royalsocietypublishing.org/content/7/4/160300.

Carlos A. Díaz-Balzac, María I. Lazaro-Peña, Lionel D. Vázquez-Figueroa, Roberto J. Díaz-Balzac, and José E. Garćia-Arrarás. Holothurian nervous system diversity revealed by neuroanatomical analysis. PLOS ONE, 11(3):1–22, 03 2016. doi:10.1371/journal.pone.0151129. URL https://doi.org/10.1371/journal.pone.0151129.

L. Díaz-Miranda, R. E. Blanco, and J. E. García-Arrarás. Localization of the heptapeptide GFSKLYFamide in the sea cucumber Holothuria glaberrima (Echinodermata): a light and electron microscopic study. J Comp Neurol, 352(4):626–640, Feb 1995. doi:10.1002/cne. 903520410. URL http://dx.doi.org/10.1002/cne.903520410.

Sarah Garner, Ivona Zysk, Glynis Byrne, Marabeth Kramer, Daniel Moller, Valerie Taylor, and Robert D. Burke. Neurogenesis in sea urchin embryos and the diversity of deuterostome neurogenic mechanisms. Development, 143(2):286–297, 2016. ISSN 0950-1991. doi:10. 1242/dev.124503. URL http://dev.biologists.org/content/143/2/286.

O Hamman. Anatomie der ophiuren und crinoiden. Ztschrft. f. Naturw. (Jena), 43:233–384, 1889.

Luke A. Hoekstra, Leonid L. Moroz, and Andreas Heyland. Novel insights into the echinoderm nervous system from histaminergic and fmrfaminergic-like cells in the sea cucumber leptosynapta clarki. PLoS One, 7(9):e44220, 2012. doi:10.1371/journal.pone.0044220. URL http://dx.doi.org/10.1371/journal.pone.0044220.

Kyle A. Long, Carlos W. Nossa, Mary A. Sewell, Nicholas H. Putnam, and Joseph F. Ryan. Low coverage sequencing of three echinoderm genomes: the brittle star ophionereis fasciata, the sea star patiriella regularis, and the sea cucumber australostichopus mollis. Giga-Science, 5(1):1–4, 2016. doi:10.1186/s13742-016-0125-6. URL +http://dx.doi.org/10.1186/s13742-016-0125-6.

Vladimir Mashanov, Olga Zueva, Tamara Rubilar, Lucia Epherra, and Jose E. García-Arrarás. Structure and Evolution of Invertebrate Nervous Systems, chapter 51 Echinodermata, pages 665–688. Oxford University Press, 2016.

Vladimir S. Mashanov, OlgaR. Zueva, Thomas Heinzeller, and IgorYu. Dolmatov. Ultrastructure of the circumoral nerve ring and the radial nerve cords in holothurians (Echinodermata). Zoomorphology, 125(1):27–38, 2006. ISSN 0720-213X. doi:10.1007/ s00435-005-0010-9. URL http://dx.doi.org/10.1007/s00435-005-0010-9.

Vladimir S. Mashanov, Olga R. Zueva, Thomas Heinzeller, Beate Aschauer, and Igor Yu. Dolmatov. Developmental origin of the adult nervous system in a holothurian: an attempt to unravel the enigma of neurogenesis in echinoderms. Evolution & Development, 9(3): 244–256, 2007. ISSN 1525-142X. doi:10.1111/j.1525-142X.2007.00157.x. URL http://dx.doi.org/10.1111/j.1525-142X.2007.00157.x.

Vladimir S Mashanov, Olga R Zueva, Thomas Heinzeller, Beate Aschauer, Wilfried W Naumann, Jesus M Grondona, Manuel Cifuentes, and Jose E Garcia-Arraras. The central nervous system of sea cucumbers (echinodermata: Holothuroidea) shows positive immunostaining for a chordate glial secretion. Front Zool, 6:11, 2009. doi:10.1186/1742-9994-6-11. URL http://dx.doi.org/10.1186/1742-9994-6-11.

Vladimir S Mashanov, Olga R Zueva, and Jose E Garcia-Arraras. Organization of glial cells in the adult sea cucumber central nervous system. Glia, 58(13):1581–1593, Oct 2010. doi:10.1002/glia.21031. URL http://dx.doi.org/10.1002/glia.21031.

Vladimir S Mashanov, Olga R Zueva, and José E García-Arrarás. Radial glial cells play a key role in echinoderm neural regeneration. BMC Biology, 11(1):49, 2013. ISSN 1741-7007. URL http://www.biomedcentral.com/1741-7007/11/49.

Vladimir S. Mashanov, Olga R. Zueva, and José E. García-Arrarás. Heterogeneous generation of new cells in the adult echinoderm nervous system. Front Neuroanat, 9:123, 2015a. doi:10.3389/fnana.2015.00123. URL http://dx.doi.org/10.3389/fnana.2015.00123.

Vladimir S. Mashanov, Olga R. Zueva, and José E. García-Arrarás. Myc regulates programmed cell death and radial glia dedifferentiation after neural injury in an echinoderm. BMC Dev Biol, 15(1):24, 2015b. doi:10.1186/s12861-015-0071-z. URL http://dx.doi.org/10.1186/s12861-015-0071-z.

Yoshiya Matsuzaka, Eiki Sato, Takeshi Kano, Hitoshi Aonuma, and Akio Ishiguro. Non-centralized and functionally localized nervous system of ophiuroids: evidence from topical anesthetic experiments. Biology Open, 6(4):425–438, 2017. doi:10.1242/bio.019836. URL http://bio.biologists.org/content/6/4/425.

Yoko Nakajima, Hiroyuki Kaneko, Greg Murray, and Robert D. Burke. Divergent patterns of neural development in larval echinoids and asteroids. Evolution & Development, 6(2): 95–104, 2004. ISSN 1525-142X. doi:10.1111/j.1525-142X.2004.04011.x. URL http://dx.doi.org/10.1111/j.1525-142X.2004.04011.x.

Sruthi Purushothaman, Sandeep Saxena, Vuppalapaty Meghah, Cherukuvada V. Brahmendra Swamy, Olga Ortega-Martinez, Sam Dupont, and Mohammed Idris. Transcriptomic and proteomic analyses of amphiura filiformis arm tissue-undergoing regeneration. Journal of Proteomics, 112:113–124, 2015. ISSN 1874-3919. doi:http://dx.doi.org/10.1016/j.jprot.2014.08.011. URL http://www.sciencedirect.com/science/article/pii/S1874391914004096.

Johannes Schindelin, Ignacio Arganda-Carreras, Erwin Frise, Verena Kaynig, Mark Longair, Tobias Pietzsch, Stephan Preibisch, Curtis Rueden, Stephan Saalfeld, Benjamin Schmid, Jean-Yves Tinevez, Daniel James White, Volker Hartenstein, Kevin Eliceiri, Pavel Tomancak, and Albert Cardona. Fiji: an open-source platform for biological-image analysis. Nat Methods, 9(7):676–682, Jul 2012. doi:10.1038/nmeth.2019. URL http://dx.doi.org/10.1038/nmeth.2019.

J. Viehweg, W. W. Naumann, and R. Olsson. Secretory radial glia in the ectoneural system of the sea star asterias rubens (echinodermata). Acta Zoologica, 79(2):119–131, 1998. ISSN 1463-6395. doi:10.1111/j.1463-6395.1998.tb01151.x. URL http://dx.doi.org/10.1111/j.1463-6395.1998.tb01151.xj.1463-6395.1998.tb01151.x.

T. Vierbuchen, A Ostermeier, Z.P. Pang, Y. Kokubu, T.C. Südhof, and M Wernig. Direct conversion of fibroblasts to functional neurons by defined factors. Nature, 463(7284):1035–41, 2010.

I. C. Wilkie. The juxtaligamental cells of ophiocomina nigra (abildgaard) (echinodermata: Ophiuroidea) and their possible role in mechano-effector function of collagenous tissue. Cell and Tissue Research, 197(3):515–530, Apr 1979. ISSN 1432-0878. doi:10.1007/BF00233574. URL https://doi.org/10.1007/BF00233574.

Iain C. Wilkie. Functional morphology of the arm spine joint and adjacent structures of the brittlestar ophiocomina nigra (echinodermata: Ophiuroidea). PLOS ONE, 11(12):1–36, 12 2016. doi:10.1371/journal.pone.0167533. URL https://doi.org/10.1371/journal.pone.0167533.

I.C. Wilkie. Mutable Collagenous Tissue: Overview and Biotechnological Perspective, pages 221–250. Springer Berlin Heidelberg, Berlin, Heidelberg, 2005. ISBN 978-3-540-27683-8. doi:10.1007/3-540-27683-1 10. URL https://doi.org/10.1007/3-540-27683-1_10.

Meet Zandawala, Ismail Moghul, Luis Alfonso Yañez Guerra, Jérome Delroisse, Nikara Abylkassimova, Andrew F. Hugall, Timothy D. O’Hara, and Maurice R. Elphick. Discovery of novel representatives of bilaterian neuropeptide families and reconstruction of neuropeptide precursor evolution in ophiuroid echinoderms. Open Biology, 7(9), 2017. doi:10.1098/rsob.170129. URL http://rsob.royalsocietypublishing.org/content/7/9/170129.

